# Evidence for biomolecular condensates of MatP in spatiotemporal regulation of the bacterial cell division cycle

**DOI:** 10.1101/2024.07.23.604758

**Authors:** Inés Barros-Medina, Miguel Ángel Robles-Ramos, Marta Sobrinos-Sanguino, Juan Román Luque-Ortega, Carlos Alfonso, William Margolin, Germán Rivas, Begoña Monterroso, Silvia Zorrilla

**Affiliations:** Department of Cellular and Molecular Biosciences, Centro de Investigaciones Biológicas Margarita Salas, Consejo Superior de Investigaciones Científicas (CSIC). 28040 Madrid, Spain; Molecular Interactions Facility, Centro de Investigaciones Biológicas Margarita Salas, Consejo Superior de Investigaciones Científicas (CSIC). 28040 Madrid, Spain; Department of Microbiology and Molecular Genetics, McGovern Medical School, UTHealth-Houston, Houston, TX 77030, USA; Department of Crystallography and Structural Biology, Instituto de Química Física Blas Cabrera, Consejo Superior de Investigaciones Científicas (CSIC). 28006 Madrid, Spain

## Abstract

An increasing number of proteins involved in bacterial cell cycle events have been recently shown to undergo phase separation. The resulting biomolecular condensates play an important role in cell cycle protein function and may be involved in development of persister cells tolerant to antibiotics. Here we report that the *E. coli* chromosomal Ter macrodomain organizer MatP, a division site selection protein implicated in the coordination of chromosome segregation with cell division, forms biomolecular condensates in cytomimetic systems. These condensates are favored by crowding and preferentially localize at the membrane of microfluidics droplets, a behavior probably mediated by MatP-lipid binding. Condensates are negatively regulated and partially dislodged from the membrane by DNA sequences recognized by MatP (*matS*), which partition into them. Unexpectedly, MatP condensation is enhanced by FtsZ, a core component of the division machinery previously described to undergo phase separation. Our biophysical analyses uncover a direct interaction between the two proteins, disrupted by *matS* sequences. This binding might have implications for FtsZ ring positioning at mid-cell by the Ter linkage, which comprises MatP and two other proteins that bridge the canonical MatP/FtsZ interaction. FtsZ/MatP condensates interconvert with bundles in response to GTP addition, providing additional levels of regulation. Consistent with discrete foci reported in cells, MatP biomolecular condensates may facilitate MatP’s role in chromosome organization and spatiotemporal regulation of cytokinesis and DNA segregation. Moreover, sequestration of MatP in these membraneless compartments, with or without FtsZ, could promote cell entry into dormant states that are able to survive antibiotic treatments.

## INTRODUCTION

Biomolecular condensation through phase separation is established as a mechanism to regulate cellular function and pathology in eukaryotes (1–3). Although its identification in bacteria is more recent, increasing evidence supports both its key role in providing a mechanism for compartmentalization of these cytoplasms lacking membrane-bound organelles, and additional levels of regulation for multiple cellular processes (4, 5). Biomolecular condensates are dynamic assemblies of a single or various scaffold biomolecules in which others, termed clients, selectively concentrate, providing spatial control to reactions (6, 7). These condensates reversibly form under a narrow range of conditions (8), being favored by multivalency and macromolecular crowding, through volume exclusion or other nonspecific weak interactions (1, 9). Proteins from *Escherichia coli*, *Bacillus subtilis* or *Caulobacter crescentus* involved in chromosome replication, segregation and cell division assemble biomolecular condensates relevant for their function, in cytomimetic systems mimicking the crowded bacterial cytoplasm or *in vivo* ((5) and references therein). In many instances, biomolecular condensation of proteins, often regulated by nucleic acids, enhances cell fitness and survival in response to different kinds of stresses. In the particular case of bacteria, condensates conferring resistance to starvation, heat shock, phage infection, oxidative stress, or treatments with antibiotics have been identified (5, 10), suggesting that these structures may represent attractive new targets for the fight of antimicrobial resistance.

MatP is a specific DNA binding protein targeting *matS* sites, which are repeated over 20 times at the Ter region of the chromosome, playing a pivotal role in its organization and compaction (11, 12). *In vivo*, MatP accumulates at precise locations in the cell, forming discrete foci (11, 13–17). Besides the specific recognition of DNA, the modular structure of MatP (13) comprises multiple regions of interaction with itself (leading to dimerization), with other proteins and with lipid membranes. Together with ZapA and ZapB, MatP forms part of the Ter-linkage that, in coordination with the Min system and SlmA-mediated nucleoid occlusion, positions the FtsZ cell division ring at mid-cell (18). Within the Ter-linkage, MatP directly interacts through its C-terminal region with ZapB (14), and ZapA links ZapB with FtsZ (19). Aside from this canonical sequence of interactions, it has been suggested that ZapB may directly bind FtsZ as well (20, 21). Through the Ter-linkage, MatP is a key player in the coordination of chromosome segregation with cell division (14). In addition, MatP interacts with the structural maintenance of chromosomes complex MukBEF, contributing to normal chromosome organization and timely unlinking of replicated chromosomes, allowing their segregation (22, 23). Thus, MatP delays sister Ter segregation by releasing MukBEF from the Ter region, which limits the availability at this location of Topoisomerase IV that normally removes chromosomal catenanes. MatP binds lipid membranes, which may play a role in the regulation of its function through modulation of its localization and recognition of other ligands (15).

One of the bacterial proteins that undergoes phase separation is FtsZ, which forms biomolecular condensates strongly promoted by macromolecular crowding (24) and through interaction with the nucleoprotein complexes of SlmA (25). FtsZ condensates and polymers interconvert in response to GTP addition and depletion after GTP hydrolysis by FtsZ, and modulation of this switch by agonists/antagonists has been proposed to contribute to the exquisite regulation of FtsZ ring assembly under normal growth conditions (26). Additionally, since they sequester FtsZ together with a strong polymerization antagonist (SlmA), these cell division condensates could be among those assembled in dormant cell states tolerant to antibiotics, in which GTP levels are reduced and vital processes like division shut down (27, 28). Consistent with this, aggresomes enriched in FtsZ have been identified in persister cells surviving antibiotic treatment that dissociate when the cells resume growth (29).

Here, by reconstitution in crowding conditions and in cell-like systems generated by microfluidics, we show that MatP forms dynamic reversible biomolecular condensates, as revealed by confocal microscopy and turbidity analyses. Condensation is strongly enhanced by FtsZ, with the two proteins assembling heterotypic condensates under conditions in which they do not significantly phase separate on their own. An orthogonal approach combining different biophysical techniques (fluorescence correlation spectroscopy, FCS; fluorescence anisotropy and sedimentation velocity, SV) uncovered direct interaction between MatP and FtsZ oligomers and GTP-triggered polymers. We also investigated the impact of *matS* sequences on the assembly of MatP biomolecular condensates and on the interactions with FtsZ. The formation of *matS-*responsive MatP biomolecular condensates, also sensitive to GTP when coassembled with FtsZ, could be relevant for the modulation of MatP spatial distribution and interaction with protein partners, to orchestrate its role in FtsZ ring positioning and chromosome organization/segregation. MatP condensates may also contribute to mechanisms determining bacterial resistance to stress conditions, as sequestration of MatP and FtsZ would promote cell cycle arrest, facilitating entrance into a stress-resistant quiescent state.

## RESULTS

### MatP forms biomolecular condensates in crowded cytomimetic systems

Confocal images of purified MatP, with Alexa Fluor 488 labeled MatP (MatP-Alexa 488) as tracer, in solutions with the crowding agent dextran revealed the presence of relatively small round structures compatible with biomolecular condensates (Figures 1a and S1a). They were also observed in the presence of Ficoll (Figure S1b), but not without crowder (Figure S1c). This was consistent with the significant turbidity of MatP solutions with crowders, and the negligible values in dilute solution (Figure 1b). To get some insight into the nature of these MatP structures, we analyzed their response to changes in ionic strength. We found that increasing salt disfavored MatP condensation, reflected in gradually lower turbidity and reduction in the abundance of condensates in the images (Figures 1c and S2a), suggesting a role of electrostatic interactions in the assembly of MatP condensates.

**Figure 1.**
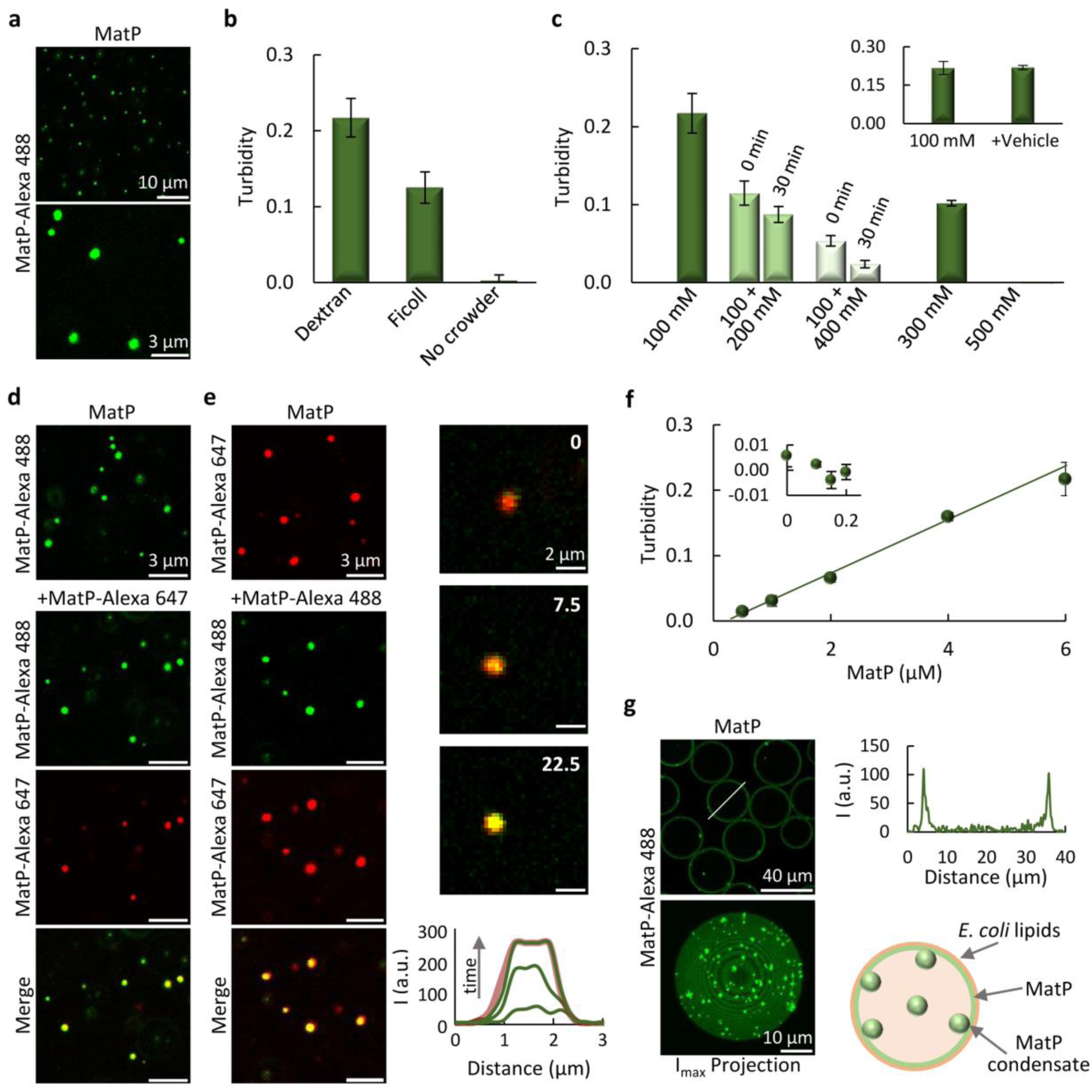
MatP forms biomolecular condensates in crowding conditions. **(a)** Confocal microscopy images of MatP condensates. **(b)** Turbidity of MatP samples in dextran (*n* > 5), Ficoll (*n* = 3) or in diluted solution (*n* = 4). **(c)** MatP turbidity decrease with the sequential addition of KCl, from 100 (*n* > 5) to 300 and 500 mM monitored at the specified times (*n* = 3), showing reversibility of MatP condensates. Samples directly prepared at these KCl concentrations are also shown (*n* = 3). Inset, control of dilution effects by adding vehicle (*n* = 3). (**d, e**) Images showing incorporation of externally added MatP into MatP condensates, indicating that they are dynamic. Right column in (e) shows the stepwise diffusion of added MatP-Alexa 488 into the condensates at the specified times in seconds (time 0, beginning of the visualization of this particular condensate) with the corresponding intensity profiles below. The profile in the red channel at 22.5 s is shown as reference. **(f)** Turbidity increase with MatP concentration (*n* = 3 except for 6 μM, *n* > 5). Line corresponds to a linear model fit, rendering the *c*_sat_ in the main text. Inset shows data below *c*_sat_ (*n* ≥ 3). Error bars are within the data symbols in most cases. **(g)** Left, images of equatorial section (top) and maximum intensity projection (bottom) of microdroplets containing MatP condensates, located mostly at the lipid membrane. Right, intensity profile corresponding to the line drawn on the image and illustration of condensates distribution. MatP concentration was 5 µM (a, d, e, g) or 6 µM (b, c). Labeled MatP was at 1 µM, except in (d-e) that was at 0.5 µM. Errors are s.d. All experiments were performed under the *MatP-crowding conditions*, substituting Ficoll for dextran or without crowder in (b) as stated.

We also asked whether these structures were dynamic and reversible, features inherent to biomolecular condensates. Condensation was reversible, as addition of supplementary KCl to 100 mM solutions containing the assemblies reduced their amount (Figure S2b) and turbidity, which approximately matched that of samples directly prepared at that final salt concentration (Figure 1c). A modest time-dependent dissociation was observed (Figure 1c), consistent with the slightly higher abundance of condensates in images acquired at short times after KCl addition compared with those of condensates directly assembled at the final KCl (*cf*. Figures S2a and S2b). Incorporation of externally added MatP, labeled with either Alexa Fluor 647 (MatP-Alexa 647) or with Alexa Fluor 488, into condensates containing MatP labeled with a spectrally different dye confirmed they were dynamic (Figures 1d, e and S2c). MatP condensation followed the characteristic behavior associated with protein phase separation in which condensates emerge above a saturation concentration (*c_sat_*) (2, 9). A *c_sat_* of 0.2 ± 0.1 µM was retrieved from the linear increase in turbidity with MatP concentration (Figure 1f), below which negligible values consistent with a single phase were obtained (Figure 1f, inset). Collectively, these experiments indicate that MatP forms dynamic reversible biomolecular condensates driven by crowding.

We further studied the properties of MatP condensates by reconstitution in microfluidics droplets containing dextran as crowder and stabilized by a monolayer of the lipid ternary mixture found in the *E. coli* inner membrane. MatP accumulated at this boundary (Figures 1g and S3), consistent with its known tendency to interact with membranes (15). Maximum intensity projections clearly showed defined condensates that, according to the distribution in the equatorial image, were located mainly within the droplets lipid surface (Figures 1g and S3). These experiments showed that MatP forms condensates in confined cell-like systems displaying crowding and a lipid membrane, with a clear preference for the latter.

### *matS* DNA accumulates in MatP condensates, generally disfavoring their formation

Since nucleic acids are among the strongest modulators of protein phase separation (1), we tested whether the *matS* DNA sequences specifically recognized by MatP influence its condensation. Interestingly, double-stranded (ds) oligonucleotides containing a single *matS* site disfavored MatP phase separation, to a higher or lower extent depending on the protein:DNA ratio and on whether *matS* was added to the condensates or incubated with MatP during sample preparation (Figure 2a). Thus, incubation with *matS* at 5:1 MatP:*matS* molar ratio only slightly decreased turbidity regarding MatP alone, while at 2:1 ratio turbidity substantially lowered, indicating that *matS* prevented significant MatP condensation at this higher concentration (Figures 2a and S4a). Notably, addition of *matS* into preassembled MatP condensates had a lower impact, reflected in a modest turbidity decrease at 5:1 MatP:*matS* ratio, somewhat larger at 2:1 ratio (Figures 2a and S4b). Consistent with these results, confocal analysis showed lower amounts of condensates when MatP was incubated with *matS* at 5:1 molar ratio, virtually disappearing at 2:1 ratio (Figures 2b and S4c), and a significant decrease in the amount of condensates when *matS* was added to them instead, at both ratios tested (Figures 2c and S4c). Interestingly, colocalization of Alexa Fluor 647 labeled *matS* (*matS*-Alexa 647) with MatP-Alexa 488 (Figures 2b, c) evidenced the partitioning of *matS* into the condensates and suggested that MatP within them was still able to bind *matS*. Therefore, specific DNA reduced MatP condensation in a concentration-dependent manner, more efficiently when MatP/*matS* complexes were formed before MatP phase separation, and incorporated into the remaining condensates.

**Figure 2.**
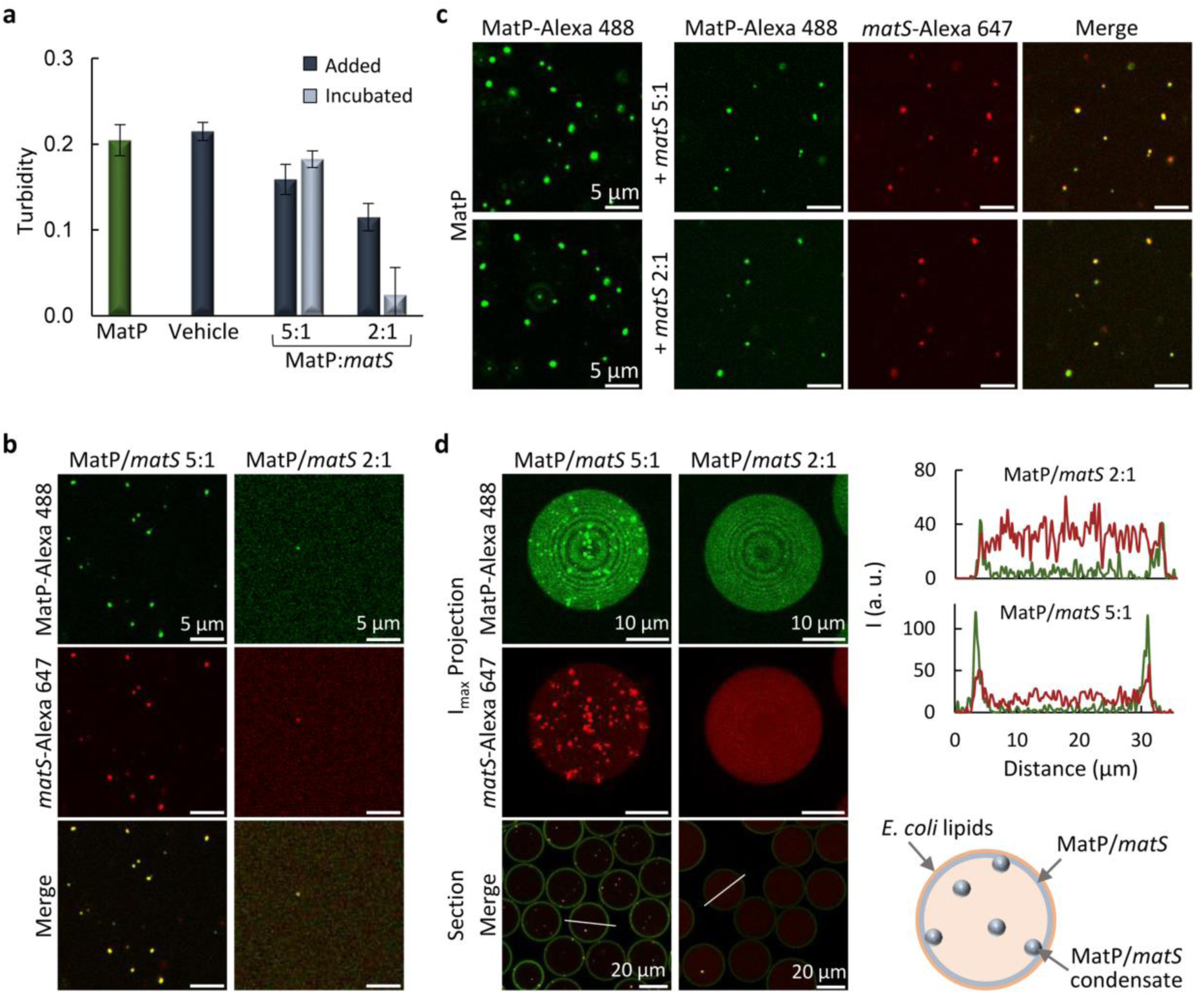
Concentration-dependent modulation of MatP condensates by *matS*. **(a)** MatP turbidity without (*n* = 5) and with *matS,* added on preformed (30 min-incubated) MatP condensates (data taken 10 min after addition, *n* ≥ 6), or incubated with MatP and the crowder for 30 min (*n* = 3). Dilution effects were ruled out by adding vehicle (*n* = 3, 10 min). Errors are s.d. (**b**, **c**) Confocal images showing concentration-dependent *matS* inhibition of MatP condensation when incubated before condensates assembly (b) or abundance reduction after addition to preformed condensates (c). MatP and *matS* colocalize in the remaining condensates. **(d)** Maximum intensity projections of the independent channels and merge equatorial sections of microdroplets containing MatP incubated with *matS*. Intensity profiles corresponding to the line drawn on the images and a scheme of condensate distribution at the lipid membrane and lumen, at 5:1 MatP:*matS* molar ratio, are shown on the right. MatP concentrations were 6 (a) or 5 µM (b, c and d) and *matS* ones corresponded to the specified MatP:*matS* molar ratios. Labeled components were at 1 µM. All experiments were performed under the *MatP-crowding conditions*.

MatP was then co-encapsulated with *matS* inside the above described microfluidics droplets (Figures 2d and S4d, e), using two aqueous streams with premixed MatP and *matS*. At 5:1 MatP:*matS* molar ratio, *matS* did not have a significant impact on the amount of condensates regarding that observed in its absence (Figure S5a), whereas at 2:1 ratio no condensates were detected (Figure 2d). The specific DNA colocalized with the MatP condensates that, still showing a preference for the membrane, increased their abundance in the lumen (Figure S5b). Intensity profiles showed accumulation of MatP at the membrane at both MatP:*matS* ratios (Figures 2d and S4d), while *matS* was distributed between lumen and membrane. These experiments showed that, when at sufficiently high concentration, *matS* hindered MatP phase separation in cell-like systems. At lower concentrations, *matS* DNA fragments partitioned into the condensates, reducing their tendency to assemble at the membrane of the microdroplets.

To evaluate the specificity of the effects of *matS* sequences, we tested ds-oligonucleotides of similar length but lacking this specific site (Figure S6). A modest reduction with respect to MatP turbidity was found when the DNA and the protein were incubated at 1:1 molar ratio prior to condensation, whereas the signal was basically unperturbed by DNA addition on preassembled condensates even after 30 min (Figures S6a and S4b). Accordingly, confocal images showed no significant variation in the condensates abundance or size under most conditions assayed (Figure S6b, c), with irregular structures only appearing upon incubation at 1:1 MatP:DNA molar ratio (Figure S6c). Sequences devoid of *matS* sites still partitioned into the condensates, probably due to nonspecific interactions with MatP favored by the high protein concentration. These results show that, unlike the specific *matS* sequences, nonspecific DNA did not significantly disrupt MatP condensates, even when it accumulated inside them.

### MatP forms heterotypic condensates with FtsZ

FtsZ has been shown to assemble heterotypic condensates in the absence of GTP, through interaction with other bacterial division proteins, acting as scaffolds (25) for additional client proteins (26). Therefore, even though FtsZ is thought to interact with MatP only indirectly (19), we asked whether there could be an interplay between FtsZ and the MatP condensates. Interestingly, FtsZ notably increased MatP solutions turbidity (Figure S7a) and confocal imaging showed numerous structures (Figure S7b, left) where the two proteins mostly colocalized, appearing more irregular than the typical round biomolecular condensates in *MatP-crowding conditions* (200 g/L dextran, 100 mM KCl, 5 mM MgCl_2_). Since biomolecular condensates usually form under a narrow range of conditions, and deviation from them often results in either absence of condensation or formation of other types of structures, we explored FtsZ/MatP mixtures at different KCl, MgCl_2_ and crowder concentrations. Reducing dextran from 200 to 150 g/L (Figure S7b) and, even more, increasing KCl to 200 mM, rendered relatively abundant structures, mostly round and hence compatible with biomolecular condensates, at both 1 and 5 mM MgCl_2_ (Figures 3a and S8).

**Figure 3.**
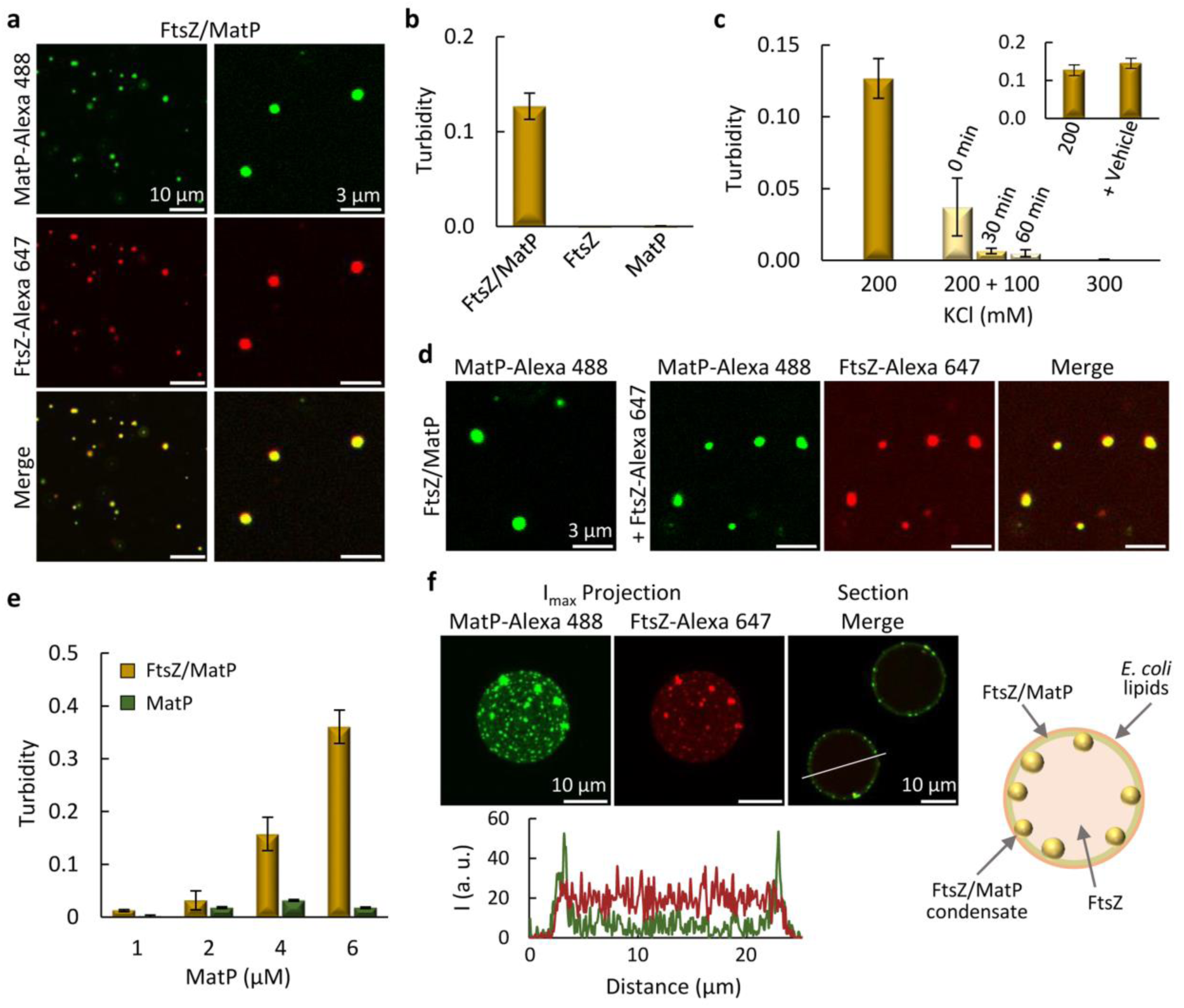
MatP forms heterotypic condensates with FtsZ. **(a)** Confocal images showing FtsZ/MatP condensates. **(b)** Turbidity of FtsZ/MatP condensates and lack of signal for only FtsZ or MatP, indicative of no condensation. **(c)** Time-dependent turbidity decrease of FtsZ/MatP condensates shifted from 200 to 300 mM KCl, indicating reversibility. Turbidity of samples prepared at 300 mM KCl is also shown. Inset reflects the absence of dilution effects, by adding vehicle. **(d)** FtsZ/MatP condensates incorporate externally-added FtsZ, stating they are dynamic. **(e)** Turbidity increase of FtsZ/MatP condensates with protein concentration, keeping a 2:1 (FtsZ:MatP) molar ratio. Turbidity of MatP alone is also shown. **(f)** Maximum intensity projections of the independent channels and merge equatorial sections of microdroplets containing condensates formed by mixing FtsZ and MatP at the droplet formation junction. On the right, schematic illustration of condensates distribution mainly at the lipid membrane and, below, intensity profiles of the green and red channels along the line depicted on the image. Concentrations were 5 µM FtsZ, 3 µM MatP and 1 µM labeled components, unless otherwise stated. Data in (b, c) are the average of 3 independent experiments, except for FtsZ/MatP at 200 mM KCl (*n* > 5); in (e) *n* = 4 and *n* = 3 for FtsZ/MatP and MatP, respectively, except when MatP was at 6 µM, *n* > 5. Errors are s.d. All experiments were performed under the *FtsZ/MatP-crowding conditions*, except for salt variations (c).

After this screening, we chose to characterize FtsZ/MatP condensates at 150 g/L dextran, 200 mM KCl and 1 mM MgCl_2_ (*FtsZ/MatP-crowding conditions*, Figure 3a). Under these conditions, no condensates were observed with FtsZ alone, in good agreement with previous reports (24), while MatP condensation was negligible (Figure S9), and significant turbidity indicative of condensation was only obtained when both proteins were present (Figure 3b). Increasing the KCl content decreased the turbidity to negligible values (Figure 3c), and drastically reduced the number of condensates in the images (Figure S10a). FtsZ/MatP condensates were also formed in the presence of Ficoll (Figure S11a), while they did not assemble without a crowding agent (Figure S11b). We next tested whether the FtsZ/MatP structures were dynamic and reversible. Addition of KCl to already assembled condensates reduced their abundance in the images and the turbidity to values similar to those of condensates formed at the final KCl concentration (Figures 3c and S10b), indicating that they were reversible. FtsZ/MatP condensates assembled with MatP-Alexa 488 as tracer readily incorporated externally added FtsZ labeled with Alexa Fluor 647 (FtsZ-Alexa 647), proving they were dynamic (Figure 3d), as further confirmed by adding MatP-Alexa 488 on preassembled FtsZ-Alexa 647 labeled condensates (Figure S10c). Turbidity measurements at a fixed 2:1 FtsZ:MatP ratio, while increasing the overall concentration, rendered negligible values at low concentrations (as for MatP solutions without FtsZ at all concentrations tested), followed by the typical concentration-dependent increase compatible with phase separation at higher concentrations (Figure 3e). Altogether, these experiments evidenced FtsZ/MatP heterotypic biomolecular condensation favored by crowding and electrostatic forces. Here, both proteins behave as scaffolds, as they both are required for condensation under conditions precluding their individual phase separation, observed under other conditions (see above and (24)).

FtsZ/MatP condensates were also analyzed in the cytomimetic platforms displaying crowding, confinement and a lipid membrane. To this end, MatP and FtsZ, with MatP-Alexa 488 and FtsZ-Alexa 647 as tracers, were jointly encapsulated inside microdroplets generated by microfluidics with dextran as crowder. Numerous condensates in which both proteins colocalized were observed almost exclusively at the membrane (Figures 3f and S12a). As in the absence of MatP, FtsZ outside the condensates was found homogeneously distributed in the lumen, and also at the membrane (*cf.* Figures 3f and S12b, and intensity profiles therein), whereas MatP was mostly at the membrane, either in the form of condensates or as a continuum, with low intensity detected in the lumen. Analogous results were obtained when encapsulating the preformed FtsZ/MatP condensates or when solutions containing each of the proteins were mixed at the droplet formation junction, to trigger their assembly just before encapsulation (*cf*. Figures 3f and S12c). These experiments showed that FtsZ and MatP form condensates in confined and crowded cell-like systems, with high preference for the lipid membrane.

### Negative regulation of condensation by *matS* is sustained for FtsZ/MatP condensates

We next asked whether the *matS* sequences specifically bound by MatP modulated the formation of FtsZ/MatP biomolecular condensates, in the same way or differently than that observed with the MatP homotypic ones. We found that, at 5:3:1 FtsZ:MatP:*matS* molar ratio, FtsZ/MatP condensates were disrupted by ds-oligonucleotides harboring a *matS* site, which largely reduced the associated turbidity, more efficiently when preincubated with the two proteins (Figures 4a and S13a). Images also showed a considerable reduction in condensate abundance and size when *matS* was added over preformed condensates (Figure 4b), and basically no condensates when *matS* was included in the FtsZ/MatP solutions from the beginning (Figures 4c and S13b). No condensates were observed either upon co-encapsulation of *matS* with the two proteins, at the same molar ratio as above, in microdroplets (Figure 4d). FtsZ was distributed between the membrane and the lumen, essentially as when encapsulated on its own, i.e. in the absence of MatP and *matS* (Figure S12b), the latter localizing primarily in the lumen of the droplet. Hence, the specific DNA recognized by MatP inhibited the formation of FtsZ/MatP condensates both in bulk crowded solution and in cell-like platforms.

**Figure 4.**
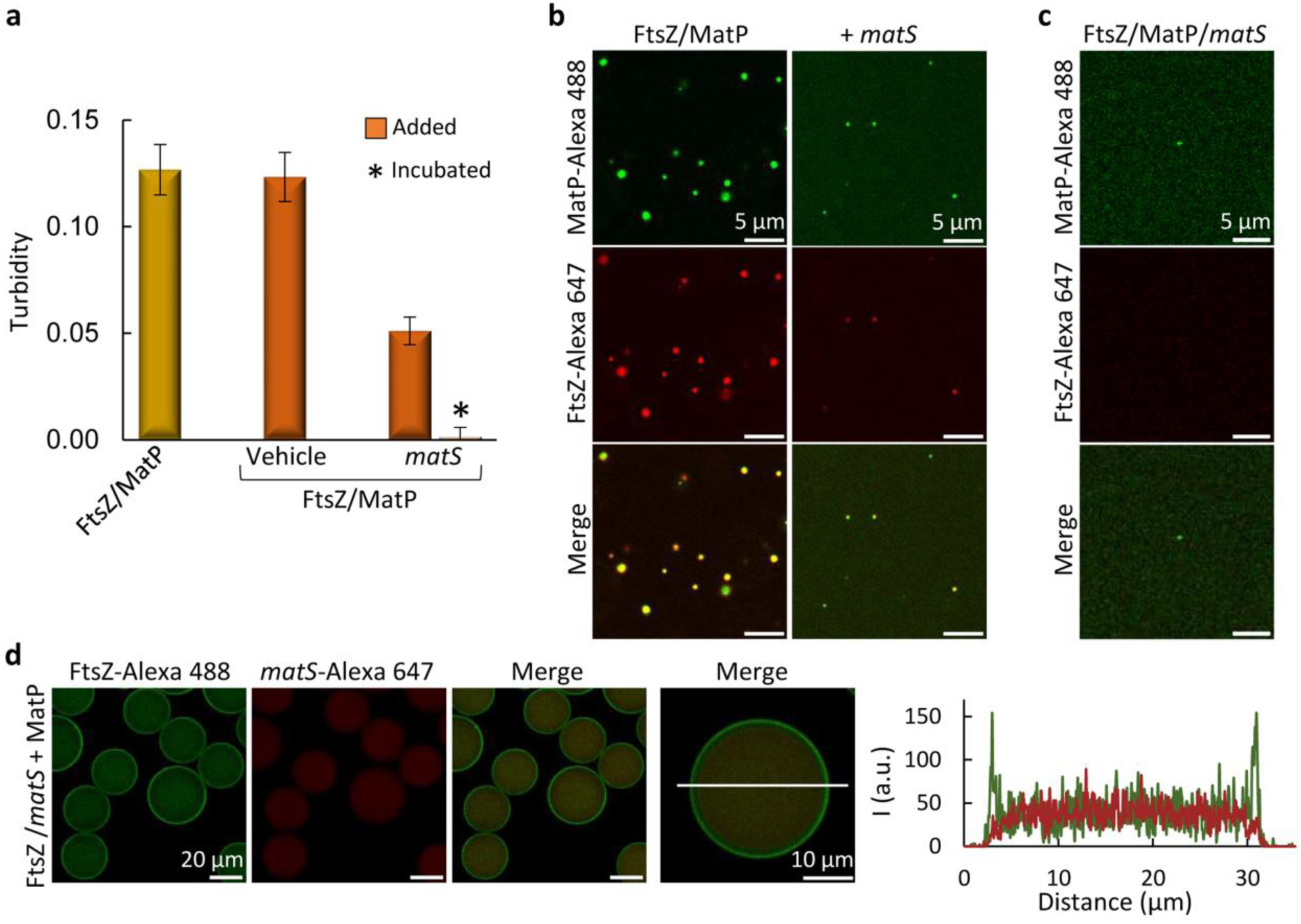
Detrimental effect of *matS* on FtsZ/MatP condensation. **(a)** Turbidity decrease of FtsZ/MatP solutions by *matS*, incubated with the protein from the beginning or added over preformed and 30 min-incubated condensates (measurements taken 10 min after addition). Dilution effects were discarded by addition of vehicle. Data are the average of three independent experiments ± s.d. (**b**, **c**) Confocal images of FtsZ/MatP condensates before and after *matS* addition (b) and when FtsZ/MatP/*matS* were incubated from the beginning (c). (**d**) Microdroplets containing FtsZ, MatP and *matS,* showing lack of condensates. FtsZ/*matS* met MatP at the droplet formation junction. On the right, intensity profiles of the green and red channels along the line drawn in the image. Concentrations were 5 µM FtsZ, 3 µM MatP and 1 µM *matS* and labeled components. All experiments were performed under the *FtsZ/MatP-crowding conditions*.

### FtsZ and MatP directly interact and *matS* inhibits the complexes

Heterotypic FtsZ/MatP biomolecular condensation under conditions at which the individual proteins cannot phase separate strongly suggests an interaction between them, which would enhance multivalency rendering the system more prone to condensation. To test this hypothesis, we probed the interaction between FtsZ and MatP in *dilute solution buffer* with 100 mM KCl by orthogonal biophysical methods.

FCS measurements of MatP, with MatP-Alexa 488 as tracer, showed autocorrelation curves (Figure 5a) described with a single diffusion coefficient *D* (69 ± 2 µm^2^/s) close to that for a MatP dimer (66 µm^2^/s), calculated from the experimentally determined sedimentation coefficient, using Svedberg equation (s = 2.5 S, (15), and see below) and the theoretical mass (∼35 kDa). Shifting of profiles to longer timescales upon addition of unlabeled FtsZ (Figure 5a) was compatible with the formation of large FtsZ/MatP complexes, with some variability in the profiles even for replicates from the same sample. Incubation of the two proteins with *matS* (at 2:1 MatP:*matS* molar ratio) rendered autocorrelation profiles superimposable with those of solely MatP (Figure 5a), meaning that *matS* inhibited the detected protein heterocomplexes. Increasing KCl to 300 mM resulted in overlapping profiles of MatP and FtsZ/MatP (Figure 5a), indicating that large (or FCS-detectable) heterocomplexes were disfavored at this higher ionic strength. Therefore, FCS analysis revealed the formation of complexes between MatP and FtsZ involving electrostatic interactions and hindered by *matS*.

**Figure 5.**
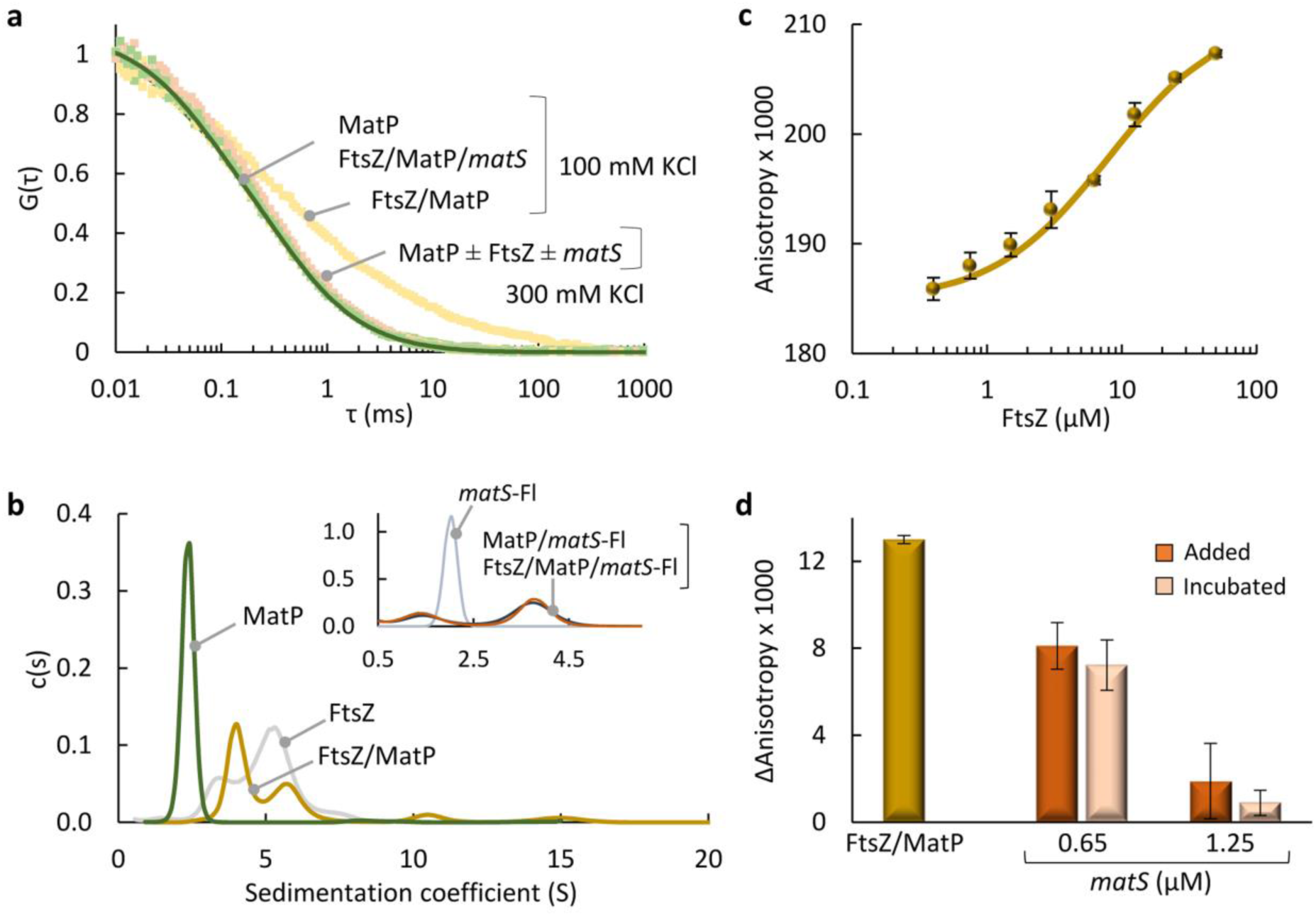
MatP binds to FtsZ through an interaction modulated by *matS*. **(a)** Normalized FCS autocorrelation curves of MatP (10 μM; 10 nM MatP-Alexa 488) and MatP with FtsZ (20 μM) in the presence and absence of *matS* (5 μM) at the specified KCl. Solid line corresponds to the fit of the model described in Materials and Methods. **(b)** SV analysis of MatP and FtsZ, alone or combined, followed at 280 nm. Inset, profiles of *matS*-Fl with MatP or FtsZ/MatP followed at 493 nm. The distribution of *matS*-Fl alone is shown for comparison. Besides species remaining in solution, FtsZ/MatP samples contained a large fraction of higher order complexes fully sedimenting before reaching the assay final speed. Concentrations were 20 μM FtsZ, 7 μM MatP and 3.5 μM *matS*-Fl. (**c)** Fluorescence anisotropy binding titrations of MatP (50 nM MatP-Alexa 488, 96 nM total MatP) with FtsZ. Solid line is the fit of the model specified in Materials and Methods and Supplementary Material. **(d)** Fluorescence anisotropy change of FtsZ (5 μM, 50 nM FtsZ-Alexa 488) in the presence of MatP (2.5 μM) with and without *matS,* either incubated for 30 min with FtsZ/MatP or added over the preincubated proteins. Data in (c, d) are the average of three independent replicates ± s.d. All experiments were performed in *dilute solution buffer* with 100 mM KCl, unless otherwise stated.

The interaction between both proteins and the effect of *matS* on the complexes were further analyzed by SV. As mentioned, MatP sedimented as a dimer (2.5 S; Figure 5b and (15)), while FtsZ solutions contained a mixture of species (Figure 5b) consistent with its tendency to form oligomers of variable size (30). Sedimentation coefficient distributions of samples containing both proteins showed the complete disappearance of the peak corresponding to MatP, and the presence of two peaks partially overlapping with those of FtsZ alone, with a substantially lower intensity signal in the higher sedimentation peak (Figure 5b). Multisignal analysis of profiles recorded at two wavelengths revealed that these peaks contained both MatP and FtsZ (Figure S14a). We detected a large fraction (*∼*50%) of higher order complexes that fully sedimented before reaching the final speed of the assay. The effect of *matS* on the FtsZ/MatP complexes was analyzed by including fluorescein-labeled *matS* (*matS*-Fl) in the samples. Sedimentation coefficient distributions showed a major peak at 3.8 S, corresponding to MatP:*matS* complexes as previously described (15), and a minor amount of free *matS* (inset in Figure 5b). This distribution was not modified in the presence of FtsZ (inset in Figure 5b), indicating that all MatP was still available to form complexes with *matS* and ruling out the formation of FtsZ/MatP complexes, in good agreement with FCS results. Therefore, SV analysis confirmed the interaction between MatP and FtsZ, including the formation of large complexes, and the inhibition by *matS*.

The FtsZ/MatP complexes were evaluated by fluorescence anisotropy binding titrations of MatP (containing MatP-Alexa 488 as tracer) with increasing concentration of unlabeled FtsZ. Analysis of the concentration-dependent increase in the anisotropy associated with heterocomplex formation (Figure 5c) rendered an apparent *K_d_* of 8 ± 3 µM (or 18 ± 6 µM, considering intensity changes; see Materials and Methods and Supplementary Material for analysis details), representing the concentration of FtsZ yielding half of the maximum signal reached. Anisotropy was also used to evaluate the effect of *matS* on the interaction, in this case with FtsZ-Alexa 488 as tracer. Addition of MatP increased the anisotropy indicating complex formation (Figure 5d and S14b), and incubation of FtsZ/MatP with *matS* or its addition over preformed complexes decreased the values (Figure 5d), reflecting a concentration-dependent inhibition of the FtsZ/MatP interactions. Anisotropy was insensitive to the addition of MatP to labeled FtsZ when KCl was increased to 300 mM (Figure S14b), a result compatible with a lack of interaction under these conditions. These experiments showed that *matS* was able to inhibit the formation of FtsZ/MatP complexes in solution and to dissociate those already formed.

Taken together, the orthogonal biophysical analyses proved the formation of heterocomplexes of moderate apparent affinity between FtsZ and MatP including large complexes, with electrostatic forces contributing to their formation and *matS* causing their disruption.

### FtsZ within FtsZ/MatP biomolecular condensates reversibly forms GTP-triggered filaments decorated with MatP

GTP-induced assembly and GTP hydrolysis are hallmarks of FtsZ functionality retained in its previously described homotypic and heterotypic biomolecular condensates (24–26). Analogously, GTP triggered bundles from FtsZ/MatP condensates, coexisting and interconnected with them (Figures 6a and S15, t <30 min). The bundles were initially weakly detected (t = 0), becoming visible shortly after (t = 10) in both the green (MatP-Alexa 488) and red (FtsZ-Alexa 647) channels, and signal colocalization strongly suggested that MatP incorporated into the FtsZ bundles. At this point, the number of condensates was significantly reduced while, as time passed, the polymers disbanded because of GDP accumulation due to GTP hydrolysis by FtsZ, and condensates where both proteins colocalized reassembled (Figures 6a and S15, t >30). Curiously enough, reassembled condensates seemed to be larger and less abundant than the ones from which polymerization was triggered (Figures 6a and S15, *cf*. GDP and t = 60-120 min). Incorporation of MatP to FtsZ bundles was further confirmed through addition of MatP to preformed FtsZ polymers (Figure 6b), allowing their inspection in the absence of condensates, and showing colocalization of the proteins. MatP seemed to prolong the lifetime of FtsZ polymers as, in its absence, no polymers could be detected 10 minutes after GTP addition (Figure S16).

**Figure 6.**
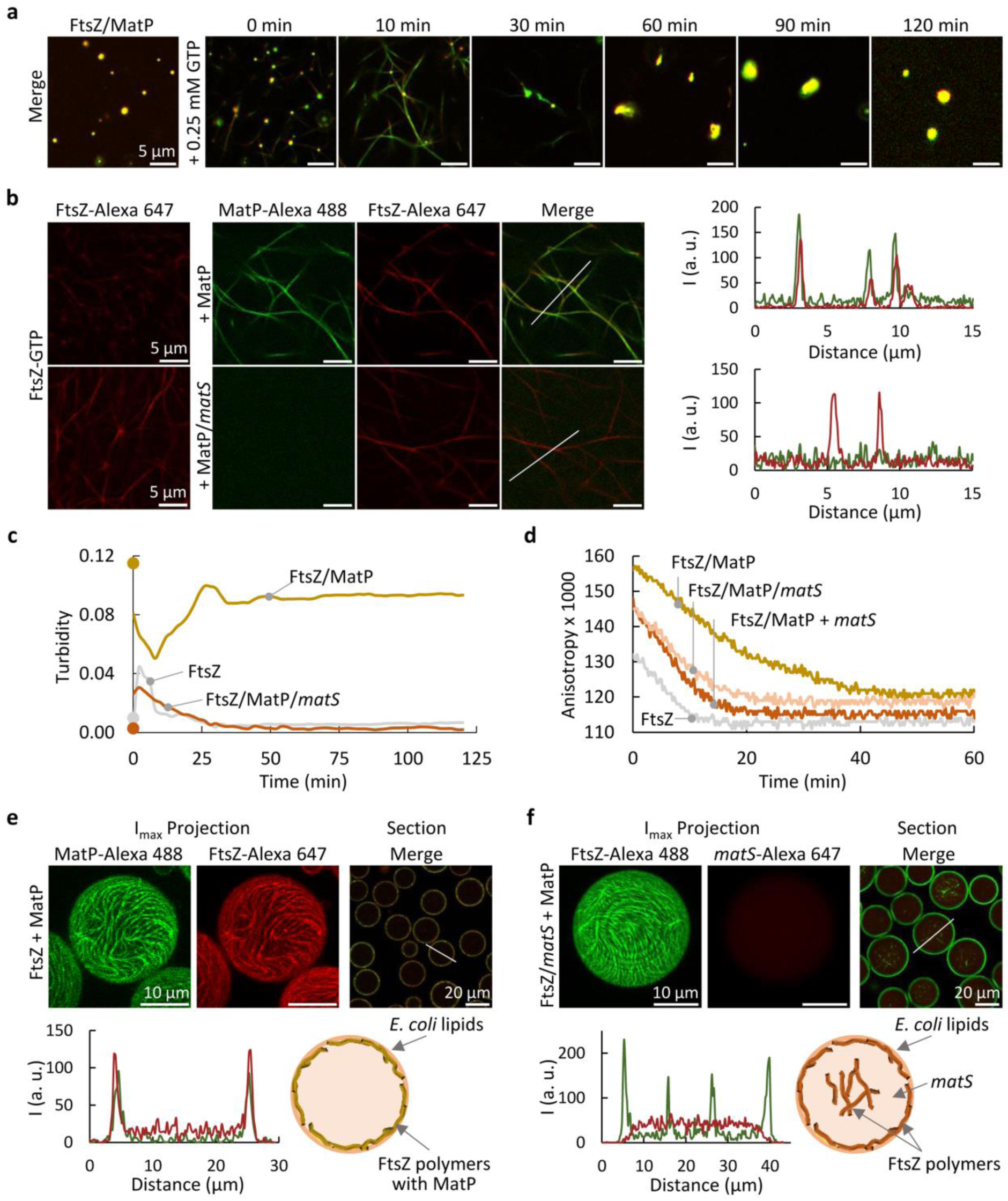
MatP decorates GTP-driven FtsZ bundles and dislodges in the presence of *matS*. **(a)** Merge confocal images of FtsZ/MatP condensates before and after triggering bundles containing FtsZ-Alexa 647 and MatP-Alexa 488 with GTP, and time-evolution showing condensate reassembly. Independent channels in Figure S15. **(b)** Images of FtsZ bundles triggered by GTP before and after addition of MatP or MatP/*matS*. On the right, intensity profiles of the red and green channels along the lines depicted in the images, evidencing FtsZ/MatP colocalization in the polymers in the absence of *matS*. **(c)** Time evolution of FtsZ-GTP turbidity as compared to FtsZ-GTP/MatP with or without *matS*. Values prior to polymerization are represented as dots at 0 min (*n* = 3). **(d)** Time evolution of fluorescence anisotropy of FtsZ-GTP (50 nM FtsZ-Alexa 488) with or without MatP or MatP/*matS* and effect of the addition of *matS* (+*matS*) to FtsZ/MatP immediately after polymerization. (**e**, **f**) Maximum intensity projections of the independent channels and merge equatorial sections of microdroplets containing FtsZ polymers and MatP without (e) and with *matS* (f). Intensity profiles of the green and red channels obtained along the line in the images are also shown. MatP met FtsZ (e) or FtsZ/*matS* (f) at the droplet formation junction. All the experiments were performed in *FtsZ/MatP-crowding conditions*. When present, the concentrations of FtsZ, MatP and *matS* were 5, 3 and 1 μM, respectively. In (a, b, e and f) concentration of labeled components was 1 μM. Polymerization was induced with 0.25 mM (a, c, d) or 2 mM GTP (b, e, f).

The response of FtsZ/MatP condensates to GTP was also monitored by turbidity (Figure 6c) and fluorescence anisotropy (Figure 6d). Turbidity decreased at short times after addition of GTP, consistent with the emergence of bundles and the concomitant dissociation of the condensates, as previously described for other FtsZ condensates (24, 26). Polymerization arising only from the protein outside the condensates would have increased the turbidity instead, because the overall signal would have been then contributed by both the intact condensates and the newly formed polymers. Turbidity reached a minimum at ∼10 min, time at which images showed mainly polymers (Figures 6a and S15), and then increased, plateauing at ∼30 minutes, (Figure 6c), a trend compatible with condensates reassembly (Figures 6a and S15, t >30 min). Without MatP, GTP induced the typical rapid increase in turbidity due to FtsZ bundles assembly, which did not form condensates, followed by a drop with polymers disassembly (Figure 6c), the low signal precluding accurate determination of how long the polymers lasted. Anisotropy showed a time-dependent decrease in the value reached upon GTP addition to FtsZ/MatP condensates (with FtsZ-Alexa 488 as tracer) due to polymers dissociation (26), plateauing after ∼40 minutes (Figure 6d), in fair agreement with confocal and turbidity analysis. Values in the presence of MatP were higher during the whole time interval than in its absence, reflecting the presence of condensates (absent from the FtsZ only samples) and/or the formation of larger or more rigid polymers. Without MatP, anisotropy plateaued at ∼10 min (Figure 6d), further supporting shorter polymer duration than with MatP. Taken together, turbidity, confocal and anisotropy showed the evolution of FtsZ/MatP condensates towards polymers in the presence of GTP and their reassembly upon GTP exhaustion, and strongly suggested that MatP binds to FtsZ bundles retarding their disassembly.

We also studied the behavior of FtsZ polymers when co-encapsulated with MatP inside microdroplets with dextran as crowder and a lipid boundary. Polymerization was triggered by GTP just before encapsulation. Confocal images showed colocalization of FtsZ-Alexa 647 and MatP-Alexa 488 tracers at the membrane (Figures 6e and S17a), forming polymers clearly observed in the maximum intensity projections and 3D reconstructions (Figure 6e and Movie 1). Without MatP, FtsZ polymers were also found at the membrane, but their presence in the lumen was more evident than with MatP (Figure S17b, and Movie 2), suggesting that the FtsZ/MatP interaction shifted them towards the membrane. Hence, MatP decorated FtsZ polymers also when reconstituted in cell-like microdroplets, increasing their tendency to locate at the lipid membrane.

### Specific DNA sequences dissociate MatP from FtsZ bundles assembled in crowding conditions

The experiments described above showed that MatP accumulated at GTP-induced FtsZ bundles formed in crowding conditions, modifying their disassembly kinetics and distribution in cytomimetic systems. We asked if *matS* could modify these observations, finding that its incubation with the proteins prevented incorporation of MatP into the polymers, its fluorescence signal appearing homogeneously distributed in the images (Figure 6b, bottom). Likewise, addition of *matS* to FtsZ polymers decorated with MatP released the latter from them (Figure S18), and MatP was not found on polymers triggered from FtsZ/MatP/*matS* samples (Figure S19). Supporting these findings, turbidity profiles depicting the evolution of FtsZ/MatP/*matS* upon GTP addition were close to those of FtsZ alone (Figure 6c), consistent with FtsZ/MatP condensates disruption (prior to GTP addition) and inhibition of MatP binding to the polymers by *matS*. Moreover, the notable difference between the anisotropy depolymerization profiles of FtsZ and FtsZ/MatP was largely reduced with *matS* either added to FtsZ/MatP polymers immediately after assembly or incubated with the proteins (Figure 6d). Therefore, *matS* precluded incorporation of MatP to FtsZ polymers and was also able to detach it when already bound.

When *matS* was included in the FtsZ/MatP mixtures encapsulated in lipid-stabilized microdroplets with crowder in their interior, GTP-triggered FtsZ polymers were observed at the membrane and also in the lumen (Figures 6f, S17c, S20 and Movie 3), adopting a distribution similar to that found in the absence of MatP (Figure S17b). MatP was detected principally at the membrane (Figure S20 and Movie 3), while *matS* remained homogeneously distributed in the lumen (Figures 6f and S17c). Neither MatP nor *matS* colocalized with FtsZ polymers (Figures 6f and S20). These experiments showed that the inhibition of MatP accumulation at FtsZ bundles by *matS* occurred also under cytomimetic conditions.

### MatP binds GTP-induced polymers of FtsZ, and specific DNA disrupts the complexes

The experiments above, performed in crowding conditions, suggested an interplay between MatP and GTP-triggered FtsZ bundles, consistent with an interaction. This prompted us to determine whether MatP could also bind the single-stranded protofilaments FtsZ forms in dilute solution (31, 32). FCS experiments with MatP-Alexa 488 as tracer showed a displacement of MatP profiles to longer times in the presence of FtsZ polymers (Figure S21a) induced by GTP and stabilized by an enzymatic regeneration system (RS, (33)), to keep them in solution during sufficient time to be characterized. SV analysis of samples containing the two proteins with GTP and RS showed the formation of very large complexes that sedimented in a large proportion before reaching the final speed of the assay, in contrast with the FtsZ polymers without MatP (Figure S21b). Finally, MatP shifted to higher values the fluorescence anisotropy of FtsZ polymers (with FtsZ-Alexa 488 as tracer, Figure S21c), a result compatible with the formation of larger and/or more rigid species. Altogether, these experiments proved MatP binding to FtsZ polymers, forming complexes larger than the FtsZ polymers without MatP.

We also tested the effect of *matS* on MatP binding to FtsZ polymers and found it disrupted the complexes detected in the FCS experiments, leading to profiles approximately matching those of samples containing only MatP, the labeled element in these assays (Figure S21a). Analogously, sedimentation coefficient distributions of FtsZ polymers were roughly the same as those observed upon incubation of FtsZ with MatP/*matS* before triggering assembly with GTP (Figure S21b). Moreover, anisotropy experiments showed a concentration-dependent antagonistic effect of *matS*, whether added prior to or after triggering polymerization of the FtsZ/MatP mixtures. Thus, the anisotropy values of FtsZ-GTP/MatP gradually approached that of FtsZ polymers without MatP as the concentration of *matS* in the samples increased (Figure S21c). Therefore, *matS* blocked MatP binding to FtsZ protofilaments in a concentration-dependent manner.

## DISCUSSION

Here, we show that the bacterial protein MatP, which participates in FtsZ ring positioning, DNA organization and coordination of cell division with chromosome segregation, undergoes biomolecular condensation in crowding cytomimetic conditions (Figures 7a and S22). These novel assemblies are negatively regulated by *matS* DNA sites that partition into them and, when formed at the microdroplets membrane for which the protein displays affinity, partially shifts the condensates towards the lumen. Accumulation of FtsZ into MatP condensates suggests a direct interaction between these two proteins, confirmed by biophysical analysis in dilute solution, that facilitates phase separation through heterotypic condensation.

**Figure 7.**
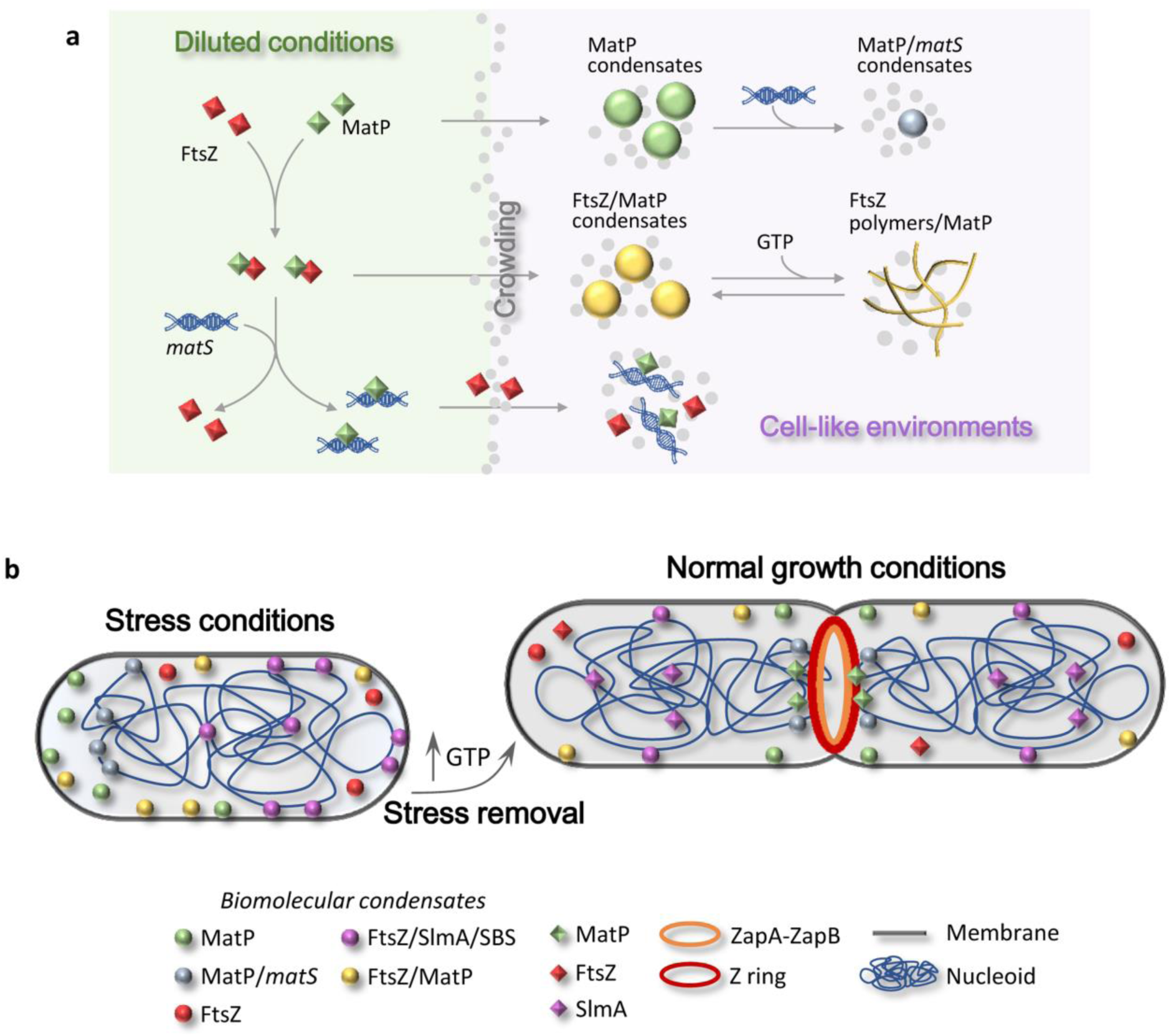
Schematic representation of the assembly and regulation of MatP biomolecular condensates and potential role under stress conditions. **(a)** Top: MatP forms crowding-driven condensates (center) where *matS* can incorporate (right). Middle: MatP interacts with FtsZ (left) and forms heterotypic condensates in crowding conditions (center) that interconvert with MatP-bound FtsZ polymers in response to GTP addition and depletion (right). Bottom: *matS* disrupts the FtsZ/MatP complexes (left), hindering condensation (right). **(b)** Under stress conditions, MatP could be transiently stored within condensates, preferentially at the membrane, and reduced GTP levels (27, 28) would favor FtsZ storage in homotypic (24) and heterotypic condensates (with MatP or SlmA (25)), which would facilitate halting MatP and FtsZ-dependent cell cycle events, to cope with stress. MatP condensates with *matS* might contribute to nucleoid compaction (a known role of MatP (11)), possibly conferring additional resistance, as described for other NAPs (5). Upon stress removal GTP increase would trigger FtsZ polymers, those arising from FtsZ/MatP condensates with bound MatP, which would be dissociated by *matS*. Some condensates may still remain at the membrane, to temporally sequester MatP from the chromosome and/or to store the large amount of FtsZ outside the division ring (32, 34) or the whole FtsZ pool under non-division conditions. MatP condensates at the membrane will be regulated by *matS* that would displace MatP to the cytoplasm/nucleoid when required.

MatP phase separation is likely driven by multivalency, as it contains multiple binding domains: a central ribbon-helix-helix mediating dimerization and interaction with *matS* sites; an N-terminal four-helix bundle also interacting with DNA; and a C-terminal flexible coiled-coil domain important for protein-protein recognition (13, 14). In this regard, recent coarse-grained molecular simulations show the presence of coiled-coil domains suffices to support biomolecular condensation (35), and many prokaryotic and eukaryotic proteins containing such domains phase separate (e.g. cell division PodJ from *C. crescentus* (36), centrosomal proteins, transcription factors and RNA binding proteins ((35) and references therein)).

MatP condensation is consistent with its originally described uneven cellular distribution pattern in bacteria (11). The protein accumulates at specific locations depending on the cell cycle stage, forming discrete foci that shift from the new cell pole to the cell center, ultimately splitting into two sister foci that remain juxtaposed until they move away from midcell. MatP foci have been reported in other *in vivo* studies, and their number and distribution analyzed under different conditions (13–17). We demonstrate here the dynamic and reversible nature of MatP biomolecular condensates and their refined control by *matS*, that could explain how the MatP pool rapidly mobilizes and reorganizes within the cell. It is possible that MatP condensates allowing subsequent incorporation of *matS* sequences, or even assembled alongside low *matS* concentrations, aid in the DNA structural arrangement mediated by this protein. The fact that these condensates are favored by macromolecular crowding would facilitate their assembly near the chromosome, which is coated by a high density of nucleoid associated proteins (NAPs), covering ∼30% of the genome in *E. coli* (37). Crowding plays a role in the phase separation of NAPs such as histone-like heat-unstable nucleoid protein (HU), DNA-binding protein from starved cells (Dps), single-stranded DNA binding protein (SSB) and RNA polymerase (RNAP) (38–41).

Interestingly, the mechanisms regulating MatP condensates are strikingly reminiscent of those reported for SSB, including sequestration of DNA at low concentrations regarding those of the proteins, and inhibition of phase separation at higher DNA concentrations (40). Furthermore, in both cases condensates assemble at the membrane with which the proteins interact. SSB condensates serve as a means to store this protein alongside interacting molecules, allowing rapid mobilization when required to repair damaged DNA. In the case of MatP, its temporal removal from the chromosome and accumulation at the membrane have been proposed to facilitate the function of proteins that help segregate the Ter region, like FtsK (15), and perhaps Topoisomerase IV (22). Condensates containing solely MatP would be detached from the membrane upon incorporation of *matS,* and high specific DNA concentrations that might locally arise would antagonize their assembly. Condensates of SSB, as well as those of Dps and HU, aggresomes, ribonucleoprotein bodies or polyP granules have been connected to stress resistance (5), conferred through DNA compaction in some instances (38, 39). MatP phase separation may have similar implications, as it is known to constrain the Ter macrodomain (42).

Our results show direct interaction between MatP and FtsZ that strongly promotes phase separation into compartments (Figures 7 and S22). These compartments could be regarded as coacervates formed by a positively and a negatively charged proteins (MatP, pI 9.5; FtsZ, pI 4.6), consistent with their strong dependence on ionic strength. FtsZ/MatP heterocomplexes were also detected in dilute conditions, including large complexes that could potentially represent incipient biomolecular condensates. These condensates could provide storage for FtsZ in non-dividing cells or accommodate the ∼60% cellular FtsZ that is not in the FtsZ ring (32, 34). MatP and FtsZ stored within these reservoirs, located principally at the membrane according to our experiments in cell-like systems, could be easily mobilized. Besides the negative regulation by *matS,* FtsZ/MatP condensates are also susceptible to disassembly by GTP, which triggers assembly of FtsZ polymers with bound MatP, ultimately dissociated from them by *matS* (Figures 7a and S22). This additional pathway is particularly interesting for preformed MatP condensates, which are more resistant to the action of *matS*. It remains to be determined whether FtsZ/MatP dissociation is required for canonical interaction of each protein with ZapA and ZapB, respectively, or if the FtsZ/MatP interaction itself is part of the Ter-linkage.

A hallmark of quiescent states is reversible arrest of cell growth and division (43), for which storage of cytokinesis factors and regulators like MatP and FtsZ inside DNA-and GTP-responsive compartments may be an effective strategy (Figure 7b), as previously proposed for FtsZ and FtsZ/SlmA condensates (24, 25). Interestingly, DNA binding strongly promotes FtsZ/SlmA phase separation (25), giving rise to condensates in which FtsZ and its inhibitor are sequestered. Such a mechanism might be expected to inhibit FtsZ ring assembly, even in the presence of agonists like ZapA (26), and would be predicted to be relevant in quiescent or dormant cells. In contrast, depending on the local concentration, *matS* binding is detrimental for FtsZ/MatP condensate formation, perhaps facilitating Ter-linkage assembly and normal cytokinesis upon stress release. The other key player in the regulation of FtsZ-containing condensates is GTP, which is strongly depleted in quiescent states (27, 28), favoring the condensation-prone oligomeric forms of FtsZ (Figure 7b). Once the stress is over, an increase in GTP concentration would disassemble these structures allowing normal cell division. Our work reinforces the idea that biomolecular condensates containing bacterial cell cycle proteins could be involved in the development of persisters and, hence, could be regarded as new targets for molecules that can kill pernicious bacteria (44).

## MATERIALS AND METHODS

Detailed information about reagents, confocal microscopy and anisotropy can be found in the Supplementary Material.

### Protein purification and labeling, and DNA hybridization

FtsZ and MatP were purified as previously described (FtsZ, calcium-induced precipitation method (30); untagged MatP (15)), and stored in aliquots at -80 °C. For the experiments, MatP was equilibrated in 50 mM Tris-HCl pH 7.5 and 300 mM KCl to avoid protein aggregation, and conditions subsequently adjusted to those conditions required for each experiment. FtsZ was dialyzed in 50 mM Tris-HCl pH 7.5, with 5 mM MgCl_2_ and the specified KCl concentration. Proteins were labeled at their amino groups with Alexa Fluor 488 or Alexa Fluor 647 carboxylic acid succinimidyl ester dyes as described (15, 45), and the labeling ratio calculated from the molar absorption coefficients of proteins and dyes (always <1 mole of dye per mole of protein). Single stranded oligonucleotides containing the *matS19* sequence recognized by MatP (11) or a nonspecific sequence of similar length, either unlabeled or labeled with Alexa Fluor 647 or fluorescein in the 5’ end, were hybridized with the complementary oligonucleotides as described (15).

### Preparation of samples in crowding bulk solution

Crowding bulk solutions were prepared in Protein LoBind® tubes (Eppendorf) by adding the protein(s) and GTP, when required, to a solution containing the specified crowder, dextran 500 or Ficoll 70, previously dialyzed in 50 mM Tris-HCl, pH 7.5 without or with 100 mM or 300 mM KCl. The final buffer conditions were adjusted to be 50 mM Tris-HCl pH 7.5, 5 mM MgCl_2_, 200 g/L dextran and 100 mM KCl (*MatP-crowding conditions*), for most analysis of MatP condensates and 50 mM Tris-HCl pH 7.5, 1 mM MgCl_2_, 150 g/L dextran and 200 mM KCl (*FtsZ/MatP-crowding conditions*), for analysis of FtsZ/MatP condensates, unless otherwise stated.

### Turbidity assays and determination of *c*_sat_

Turbidity measurements (at 350 nm) were performed using a Varioskan Flash Plate reader (Thermo Fisher Scientific, MA, USA) following a protocol described elsewhere (24) but with 85 µL samples in 384-well clear polystyrene, flat bottom microplates (Greiner Bio-One). Salt shift experiments were conducted by adding a small volume of a KCl solution into the condensates samples. Vehicle refers to controls where an equivalent volume of buffer was added instead, accounting for possible dilution effects. The concentration threshold for condensate formation, *c*_sat_, was determined by fitting a linear model to the data scaling with the protein concentration as described earlier (24, 46), and corresponds to the x intercept. Data collection started after 30 min incubation at room temperature or, in samples monitored with time, immediately after addition of the studied element (*matS*, GTP, etc.). Results are the average ± s.d. or representative profiles from at least three independent experiments.

### Microfluidics encapsulation

Encapsulation, in microfluidic devices produced by conventional soft lithography (47), involved mixing two aqueous streams in a 1:1 ratio just before the droplet formation junction. For multiple elements, aqueous phases of various compositions were assayed accounting for possible effects from incubation times: *matS*/MatP and FtsZ/MatP, including the two elements together in both aqueous phases or each one in a different solution with analogous results; FtsZ/MatP with *matS*, the latter included either in the stream with FtsZ or with MatP, with similar results. FtsZ polymerization was induced just before encapsulation, adding GTP in the FtsZ-free aqueous solution. The third stream contained the mineral oil with the *E. coli* lipid mixture (20 g/L), prepared as described earlier (15). Solutions were delivered at constant flows of 150 μL/h (oil phase) and 10 μL/h (aqueous phases) by automated syringe pumps (Cetoni GmbH, Germany), and production in the microfluidic device was monitored with an Axiovert 135 fluorescence microscope (Zeiss). Microdroplets were collected and observed 30 min after production.

### Confocal microscopy

Samples, either crowding bulk solutions or microfluidics droplets, were visualized using a Leica TCS SP5 inverted confocal microscope as described (26). Condensates were incubated for 30 min before visualization. For each sample, several images were registered. Production of images and data analysis was done with ImageJ (48). Intensity profiles, maximum intensity projections and 3D reconstructions, always corresponding to the raw images, were obtained with the straight-line, Z-projection and 3D-projection tools, respectively. Condensates diameter distribution were obtained with the particle analysis option as described (24). Brightness was uniformly increased in the whole image, only in the specified cases, using Microsoft PowerPoint. Representative images of at least three individual experiments are shown, except for microfluidic encapsulations that were repeated at least twice.

### Fluorescence correlation spectroscopy (FCS)

FCS measurements were conducted on a Microtime 200 (PicoQuant) time-resolved confocal fluorescence microscope equipped with a pulsed laser diode head (LDH-P-C-485) for excitation, as previously described (26). Experiments were done in *dilute solution buffer*: 50 mM Tris-HCl pH 7.5, 5 mM MgCl_2_ and 100 or 300 mM KCl, as specified. For FCS measurements, 0.05 g/L BSA was added, to avoid nonspecific adsorption to the surfaces, also prevented by using pegylated coverslips. Shown autocorrelation curves are representative of, at least, three independent experiments (5 profiles each). Analysis was performed with the FFS Data Processor Software (version 2.4 extended, SSTC (49)), using models with a term for the triplet state dynamics and one-diffusing species for the MatP or FtsZ/MatP/*matS* curves. Reported diffusion coefficients are the average of three independent measurements ± s.d. Quantitative analysis was omitted for FtsZ/MatP in view of the multiplicity of species present, including very large complexes.

### Sedimentation velocity assays (SV)

Samples, in *dilute solution buffer* with 100 mM KCl, were centrifuged at 48,000 rpm in an Optima XL-I analytical ultracentrifuge (Beckman-Coulter Inc.) as earlier described (26), and profiles were recorded by absorbance at 250 or 280 nm, or at 280 and 493 nm when including *matS*-Fl. When specified, GTP and the aforementioned GTP RS were included, and profiles recorded by Raleigh interference. Differential sedimentation coefficient distributions were calculated by SEDFIT (50), and heterocomplex formation was qualitatively characterized by multi-signal sedimentation velocity (MSSV) with SEDPHAT (51).

### Fluorescence anisotropy

Anisotropy experiments were performed in a Spark® Multimode microplate reader (Tecan) as described (26). FtsZ/MatP binding isotherms were obtained using 50 nM MatP-Alexa 488 as tracer, in the same buffer used for the FCS experiments, with 100 mM KCl. Binding analysis was conducted with a simple 1:1 model using BIOEQS software (52, 53), to obtain apparent *Kd*s, corresponding to the FtsZ concentration rendering half of maximum binding. Uncertainties in the parameters retrieved were calculated with the same software by rigorous confidence limit testing at the 67% level, and error propagation. The effect of *matS* on the FtsZ/MatP interaction and titrations of FtsZ with MatP were conducted in *dilute solution buffer* with 100 or 300 mM KCl, as specified. Anisotropy of FtsZ polymers with or without MatP and/or *matS* was monitored with time, after GTP addition in *FtsZ/MatP-crowding conditions* or measured at time 0 in *dilute solution buffer* with 100 mM KCl. Reported anisotropy data are the average ± s.d. or representative profiles from, at least, three independent experiments.

## DATA AVAILABILITY

All data are available from the corresponding authors upon request.

## SUPPLEMENTARY MATERIAL

Supplementary Data are available online.

## AUTHOR CONTRIBUTIONS

Inés Barros-Medina: investigation, formal analysis, visualization and writing— review and editing. Miguel Ángel Robles-Ramos: data curation, formal analysis, visualization, investigation and writing— review and editing. Marta Sobrinos-Sanguino: formal analysis, investigation and writing— review and editing. Juan Román Luque-Ortega: formal analysis and investigation. Carlos Alfonso: formal analysis and investigation. William Margolin: validation, funding acquisition, writing-review and editing. Germán Rivas: validation, funding acquisition, writing-review and editing. Begoña Monterroso: conceptualization, data curation, formal analysis, project administration, supervision, validation, writing—original draft, writing—review and editing. Silvia Zorrilla: conceptualization, data curation, formal analysis, funding acquisition, project administration, supervision, validation, writing—original draft and writing—review and editing. All authors gave final approval for publication and agreed to be held accountable for the work performed therein.

## ACKNOWLEDGEMENTS

We thank the staff of CIB Margarita Salas Confocal Laser and Multidimensional Microscopy (M.T. Seisdedos and G. Elvira) and Molecular Interactions (Ó. Nuero) Facilities for excellent technical assistance, and the Technical Support Facility for invaluable input. We also thank Drs. V. Buschmann and E. Sisamakis for generous technical help with the time-resolved confocal instrument, M. Schumacher (Duke University) for kindly providing the MatP plasmid, and W. T. S. Huck and A. Piruska (Radboud University) for the kind gift of microfluidics silicon masters.

## FUNDING

This work was supported by Grant PID2019-104544GB-I00 funded by MICIU/AEI/10.13039/501100011033 and by Grant PID2022-136951NB-I00 funded by MICIU/AEI/10.13039/501100011033 and by ERDF, EU. W.M. was supported by National Institutes of Health grant GM131705. I.B.-M. was supported by Grant FPU2020-05620 funded by MICIU/AEI/10.13039/501100011033 and by ESF Investing in your future. M.A.R.-R. was supported by Grant BES-2017-082003 funded by MICIU/AEI/10.13039/501100011033 and by ESF Investing in your future. M.S.-S. was supported by Grant PTA2020-018219-I funded by MICIU/AEI/10.13039/501100011033. The Systems Biochemistry of Bacterial Division group (CIB Margarita Salas) participates in the CSIC Conexiones LifeHUB (PIE-202120E047). The funders had no role in study design, data collection and interpretation, or the decision to submit the work for publication.

## CONFLICT OF INTEREST

We declare we have no competing interests.

## Notes

### Competing Interest Statement

The authors have declared no competing interest.

### Summary of Updates

Text has been shortened and number of figures reduced.

