## Supplementary Material for "Evidence for biomolecular condensates of MatP in spatiotemporal regulation of the bacterial cell division cycle"

[1] Department of Cellular and Molecular Biosciences, Centro de Investigaciones Biológicas Margarita Salas, Consejo Superior de Investigaciones Científicas (CSIC). 28040 Madrid, Spain
[2] Molecular Interactions Facility, Centro de Investigaciones Biológicas Margarita Salas, Consejo Superior de Investigaciones Científicas (CSIC). 28040 Madrid, Spain
[3] Department of Microbiology and Molecular Genetics, McGovern Medical School, UTHealth-Houston, Houston, TX 77030, USA
[4] Department of Crystallography and Structural Biology, Instituto de Química Física Blas Cabrera, Consejo Superior de Investigaciones Científicas (CSIC). 28006 Madrid, Spain
† Equal contribution as first authors, in alphabetical order

| Table of contents | Page | | Page |
|---|---|---|---|
| Figure S1 ................................ | s2 | Figure S17 .................................................... | s11 |
| Figure S2 ................................ | s2 | Figure S18 .................................................... | s11 |
| Figure S3 ................................ | s3 | Figure S19 .................................................... | s12 |
| Figure S4 ................................ | s3 | Figure S20 .................................................... | s12 |
| Figure S5 ................................ | s4 | Figure S21 .................................................... | s13 |
| Figure S6 ................................ | s5 | Figure S22 .................................................... | s13 |
| Figure S7 .............................. | s6 | Legends for Supplementary Movies .......... | s14 |
| Figure S8 ................................ | s6 | Supplementary Materials and Methods | |
| Figure S9 ................................ | s7 | Reagents | s14 |
| Figure S10 .............................. | s7 | Confocal microscopy | s14 |
| Figure S11 .............................. | s8 | Fluorescence anisotropy | s15 |
| Figure S12 .............................. | s8 | Supplementary references | s15 |
| Figure S13 .............................. | s9 | | |
| Figure S14 .............................. | s9 | | |
| Figure S15 .............................. | s10 | | |
| Figure S16 .............................. | s10 | | |

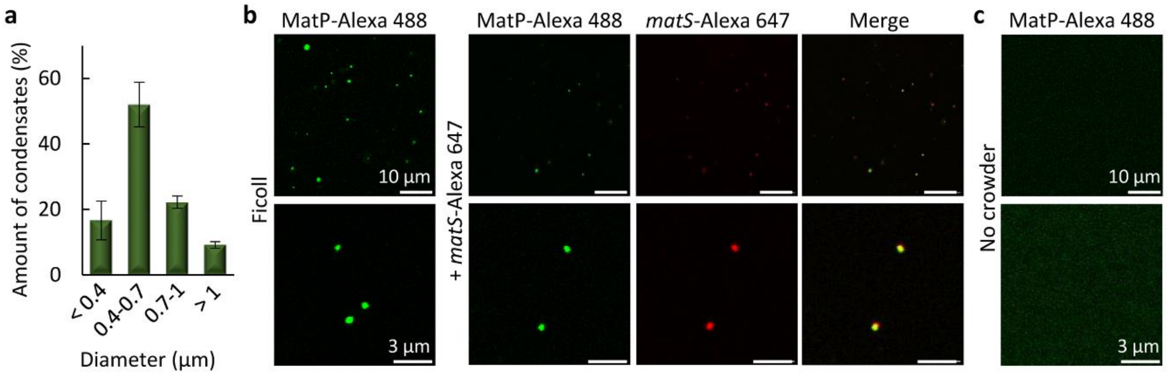


**Figure S1.** MatP condensates are formed in crowding conditions and *matS* partitions into them.
(**a**) Size distribution of MatP condensates obtained from confocal images ($n$ = 542 particles). Errors are s.d. from 6 independent images. The average number of condensates per image was approximately 90.
(**b**) Confocal images showing MatP condensates in Ficoll (200 g/L) before and after addition of *matS*.
(**c**) Images showing absence of MatP condensates in diluted solution.
Concentrations were 5 µM MatP and 1 µM labeled components. Experiments in (a, b) were performed in *MatP-crowding conditions*, except for the crowder substituted by Ficoll in (b). Experiments in (c) were conducted in *dilute solution buffer* with 100 mM KCl.

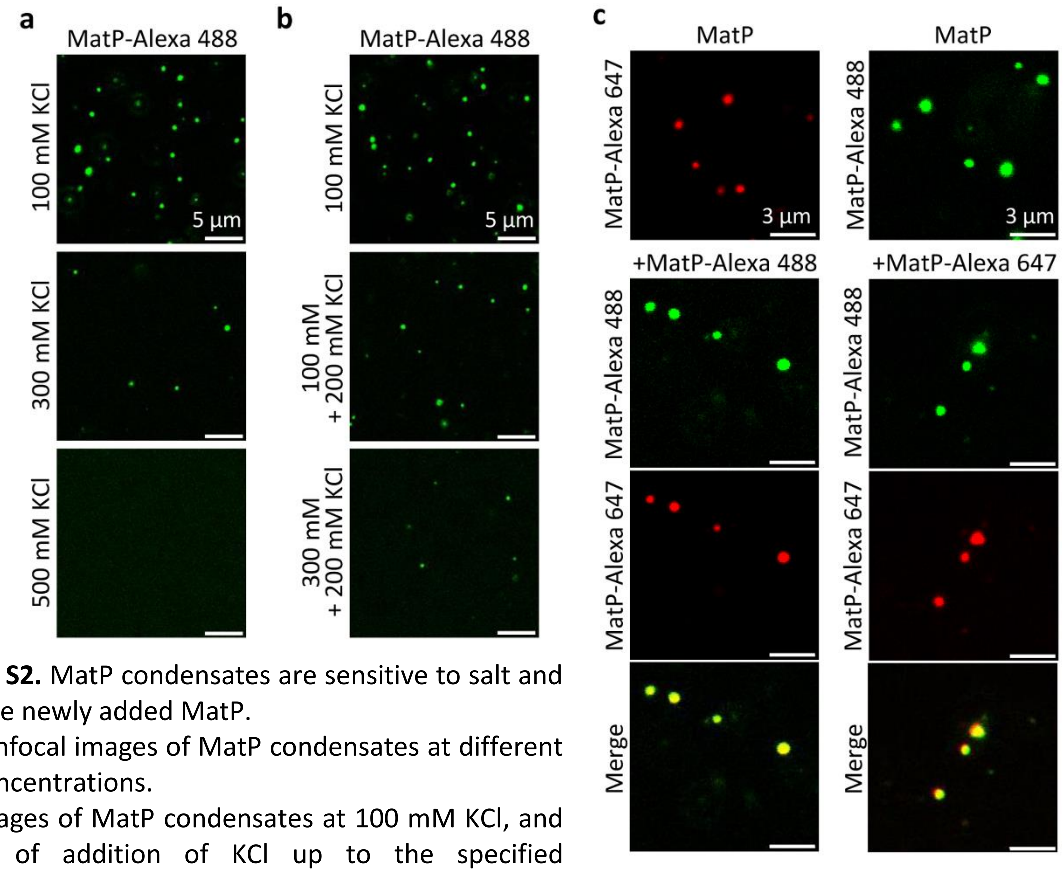


**Figure S2.** MatP condensates are sensitive to salt and capture newly added MatP.
(**a**) Confocal images of MatP condensates at different KCl concentrations.
(**b**) Images of MatP condensates at 100 mM KCl, and effect of addition of KCl up to the specified concentrations, immediately after addition. Dilution effects were ruled out in controls adding vehicle (Figure 1c).
(**c**) Images of MatP condensates (with MatP labeled with the specified dye as tracer) before and after addition of MatP labeled with a spectrally different dye.

Concentrations were 5 (a, b) or 10 μM MatP (c) and 1 (a, b) or 0.5 μM (c) labeled components. Experiments were performed in *MatP-crowding conditions* and at the specified KCl concentration in (a).

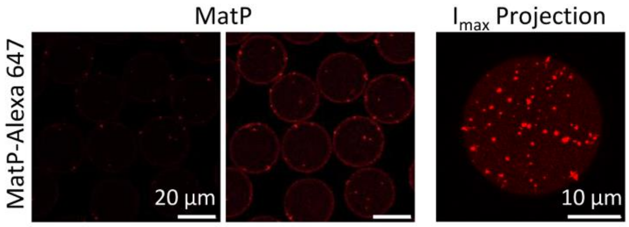


**Figure S3.** MatP forms condensates when encapsulated inside microfluidic droplets. Confocal image of equatorial sections of microdroplets containing MatP (left and middle image are the same, the latter with 60% enhanced brightness, see Materials and Methods in main text). Right, maximum intensity projection of a microdroplet. Concentration was 5 μM MatP (with 1 μM MatP-Alexa 647). Experiments were performed in *MatP-crowding conditions.*

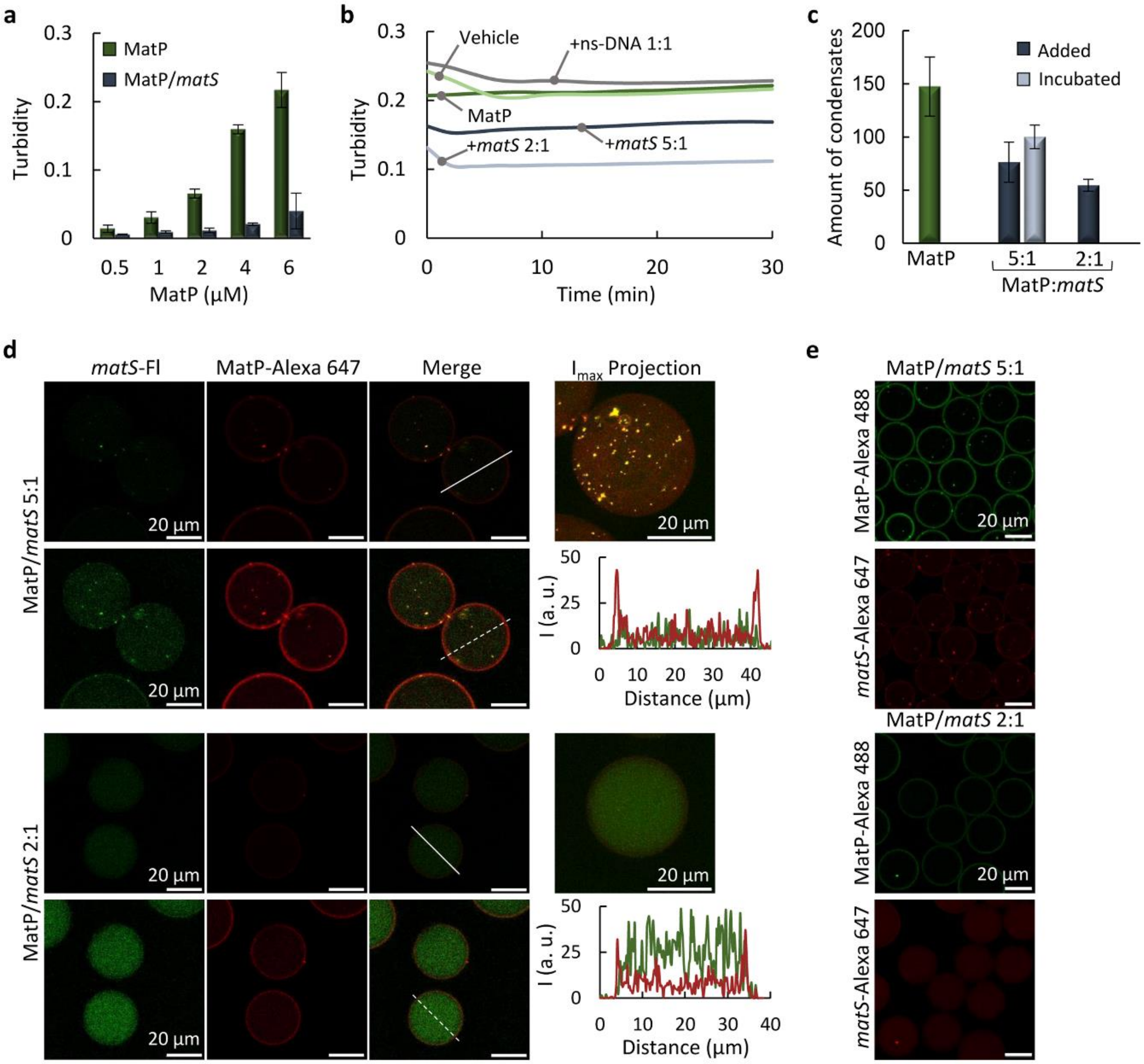


**Figure S4.** Modulation of MatP condensates by *matS*.

(**a**) Concentration dependent turbidity of MatP without and with *matS* incubated from the beginning ($n \geq 3$; MatP:*matS* maintained at a 2:1 ratio).
(**b**) Evolution with time of MatP condensates turbidity, without and with added *matS* at different ratios or nonspecific DNA (ns-DNA). Dilution effects were discarded by adding vehicle (volume equivalent to the highest used in these assays).
(**c**) Average number of condensates without and with *matS* added on preformed condensates or incubated with MatP from the beginning at the specified ratios. 9 (without *matS*) or 6 (with *matS*) independent confocal images (82 x 82 µm fields) were analyzed.
(**d**) Confocal images of equatorial sections of microdroplets containing MatP incubated with *matS* at the specified ratios. At each MatP:*matS* ratio, top and bottom row images are the same, the latter with brightness uniformly increased (60% and 55% for 5:1 and 2:1, respectively, see Materials and Methods in the main text). Intensity profiles along the line drawn in the raw images (indicated as dashed lines in the brightness-increased images to facilitate visualization) and merged maximum intensity projections are shown on the right.
(**e**) Images of equatorial sections of microdroplets containing MatP incubated with *matS* at the specified ratios and with the indicated tracers (merge and maximum intensity projections in Figure 2d).
Concentrations were 6 (b) or 5 µM MatP (c-e), and 1 µM labeled components. Indicated MatP:*matS* ratios are in molar. Errors are s.d. Experiments were performed in *MatP-crowding conditions*.

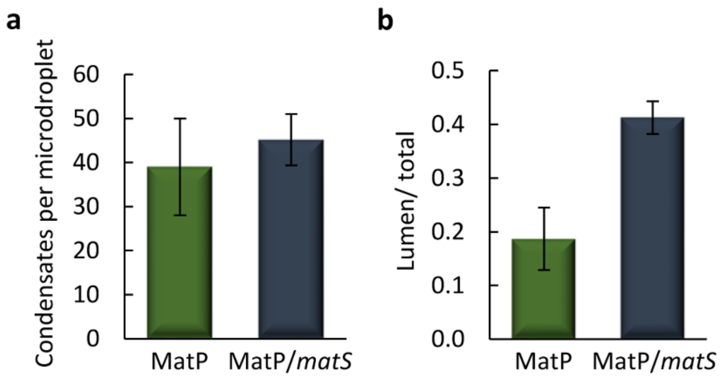


**Figure S5.** *matS* does not significantly modify the amount of condensates in microdroplets but shifts them toward the lumen.
(**a**) Average number of condensates per microdroplet, and
(**b**) fraction of condensates in the lumen of microdroplets containing MatP or MatP/*matS* in a 5:1 molar ratio.
Concentrations were 5 µM MatP and 1 µM *matS*. Experiments were performed in *MatP-crowding conditions.*

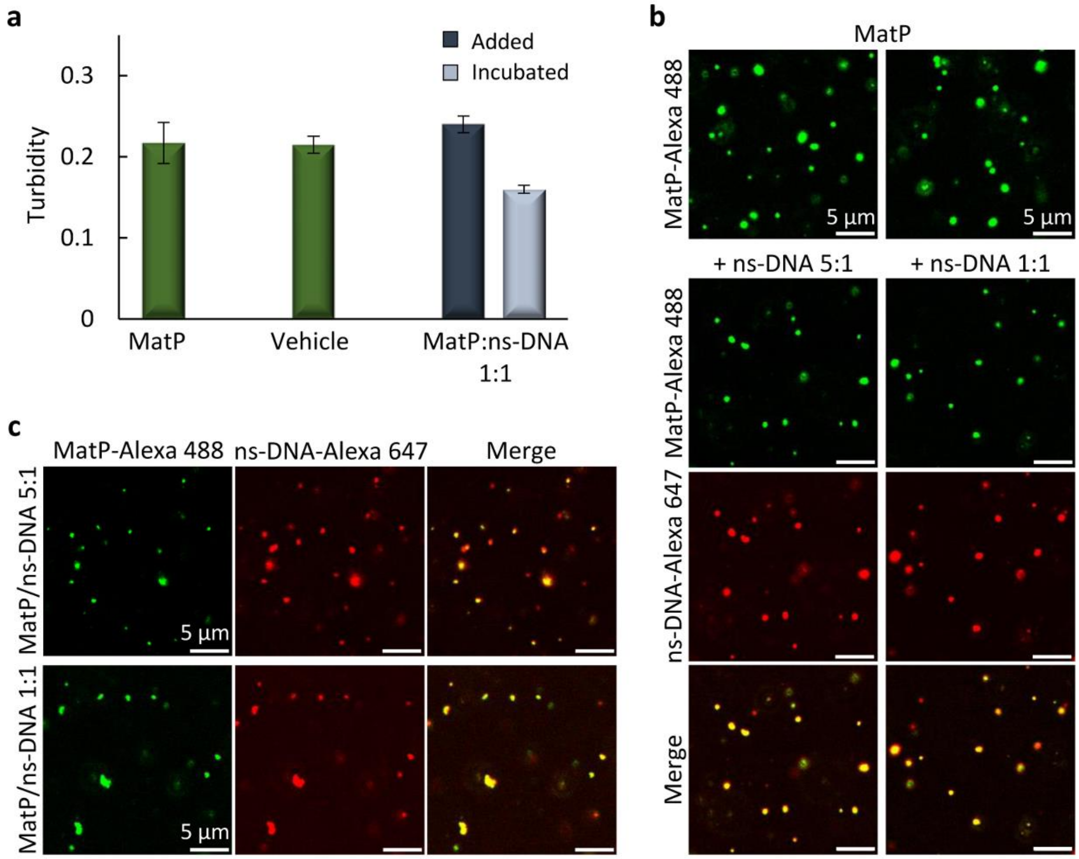


**Figure S6.** Nonspecific DNA does not significantly disrupt MatP condensates and partitions into them. (**a**) Turbidity of MatP samples without ($n > 5$) and with high concentration of nonspecific DNA (ns-DNA; $n = 3$), added or incubated. Addition of vehicle ($n = 3$, time 0) discards dilution effects. Errors are s.d.
(**b**) Confocal images showing colocalization of added ns-DNA and MatP in condensates.
(**c**) Images showing colocalization of incubated ns-DNA and MatP in condensates.
Concentrations were 5 µM MatP and 1 µM labeled components. Indicated MatP:ns-DNA ratios are in molar. Experiments were performed in *MatP-crowding conditions*.

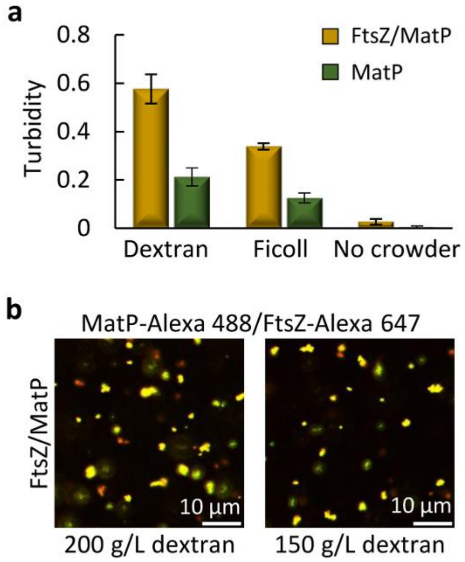


**Figure S7.** FtsZ and MatP form heterotypic condensates under crowding conditions.

(**a**) Turbidity of MatP without and with FtsZ under *MatP-crowding conditions* (i.e. with dextran as crowder; $n > 5$), or under the same conditions but replacing dextran by Ficoll ($n = 3$) or without crowder ($n = 4$). Data are the average of the indicated independent experiments ± s.d.

(**b**) Confocal images of FtsZ/MatP condensates in *MatP-crowding conditions* or under the same conditions but with 150 g/L dextran. Images are the merge of the green and red channels.

Concentrations were 12 (a) or 5 µM (b) FtsZ, 6 (a) or 3 µM (b) MatP and 1 µM labeled elements (b).

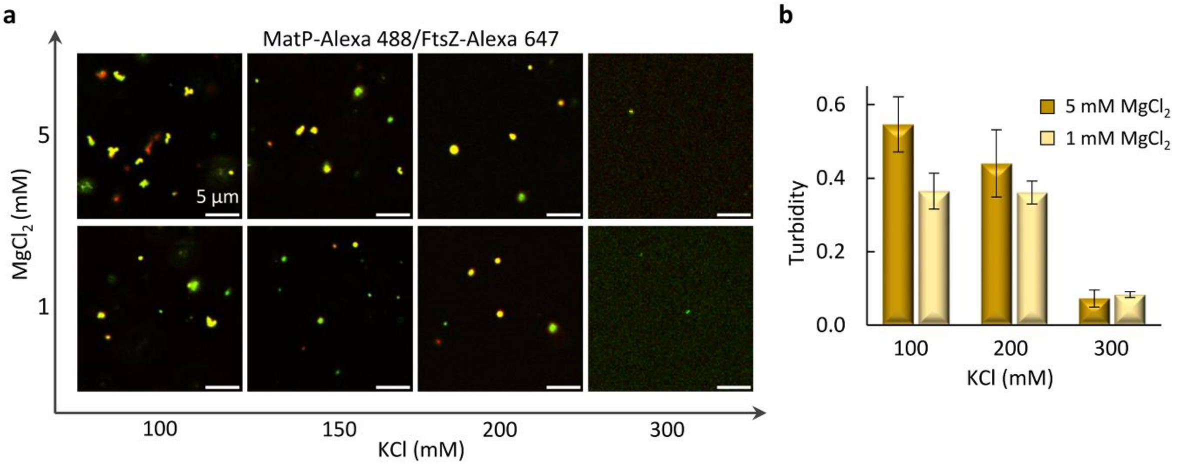


**Figure S8.** FtsZ/MatP condensates are sensitive to solution conditions.

(**a**) Merge confocal images of the green and red channels, showing the effect of KCl and $MgCl_2$ on FtsZ/MatP condensates.

(**b**) Dependence of the turbidity signal of FtsZ/MatP condensates with KCl and $MgCl_2$ concentrations. $n = 3$ for all samples but 200 mM KCl and 1 mM $MgCl_2$, $n > 5$.

FtsZ and MatP concentrations were 5 and 3 µM (a) or 12 and 6 µM (b), respectively, and labeled components (a) were at 1 µM. Experiments were performed in buffer 50 mM Tris-HCl pH 7.5, 150 g/L dextran.

**Figure S9.** Assembly of condensates by FtsZ or MatP under the *FtsZ/MatP-crowding conditions* is highly disfavored.
Absence or reduced number of condensates in samples with 5 µM FtsZ or 5 µM MatP, respectively, in *FtsZ/MatP-crowding conditions*. Concentration of labeled elements was 1 µM. Note that the concentration of MatP is higher in these experiments than in those shown in Figure 3 in the main text.

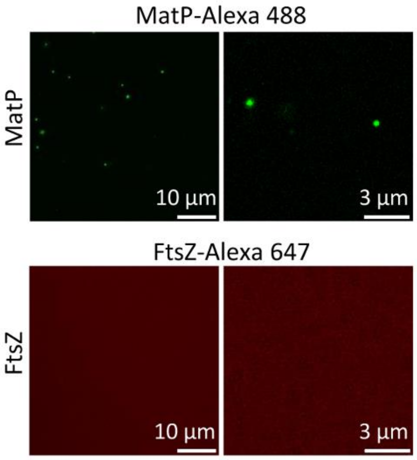


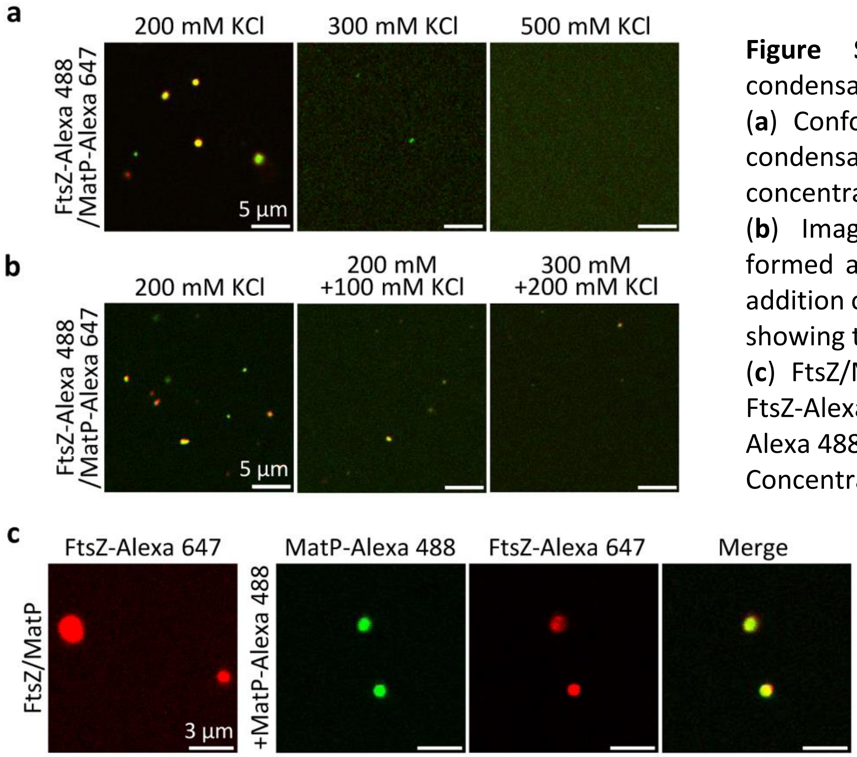


**Figure S10.** FtsZ/MatP biomolecular condensates are reversible and dynamic.
(**a**) Confocal images showing FtsZ/MatP condensates prepared at the stated salt concentrations.
(**b**) Images of FtsZ/MatP condensates formed at 200 mM KCl before and after addition of the specified KCl concentration showing their dissociation.
(**c**) FtsZ/MatP condensates labeled with FtsZ-Alexa 647 capture newly added MatP-Alexa 488.
Concentrations were 5 µM FtsZ, 3 µM MatP and 1 µM labeled components. Experiments were performed in *FtsZ/MatP-crowding conditions*, with the KCl concentration in (a) modified as stated.

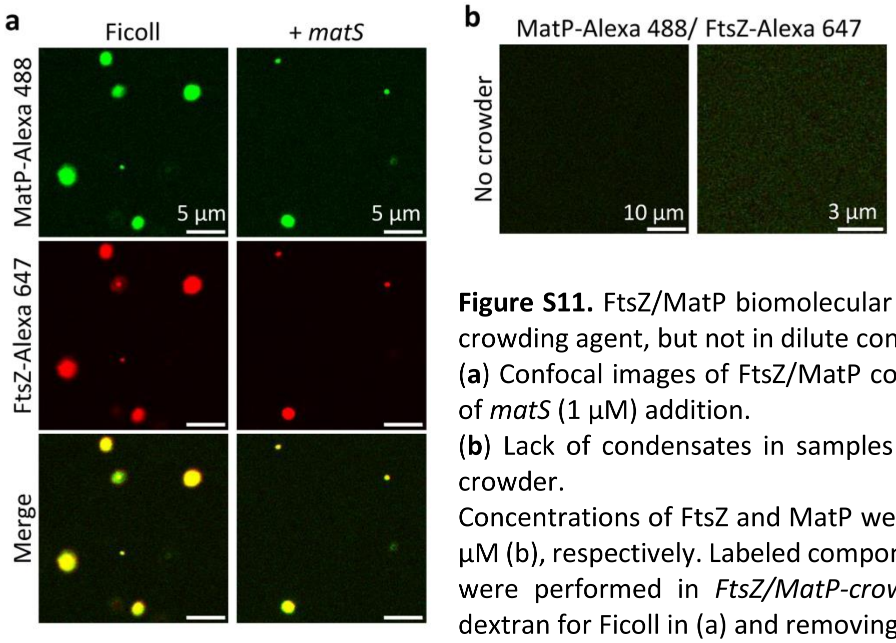


**Figure S11.** FtsZ/MatP biomolecular condensates form in Ficoll as crowding agent, but not in dilute conditions, and respond to *matS*. (**a**) Confocal images of FtsZ/MatP condensates in Ficoll and effect of *matS* (1 µM) addition.
(**b**) Lack of condensates in samples with FtsZ and MatP without crowder.
Concentrations of FtsZ and MatP were 12 and 6 µM (a) or 5 and 3 µM (b), respectively. Labeled components were 1 µM. Experiments were performed in *FtsZ/MatP-crowding conditions* substituting dextran for Ficoll in (a) and removing the crowder in (b).

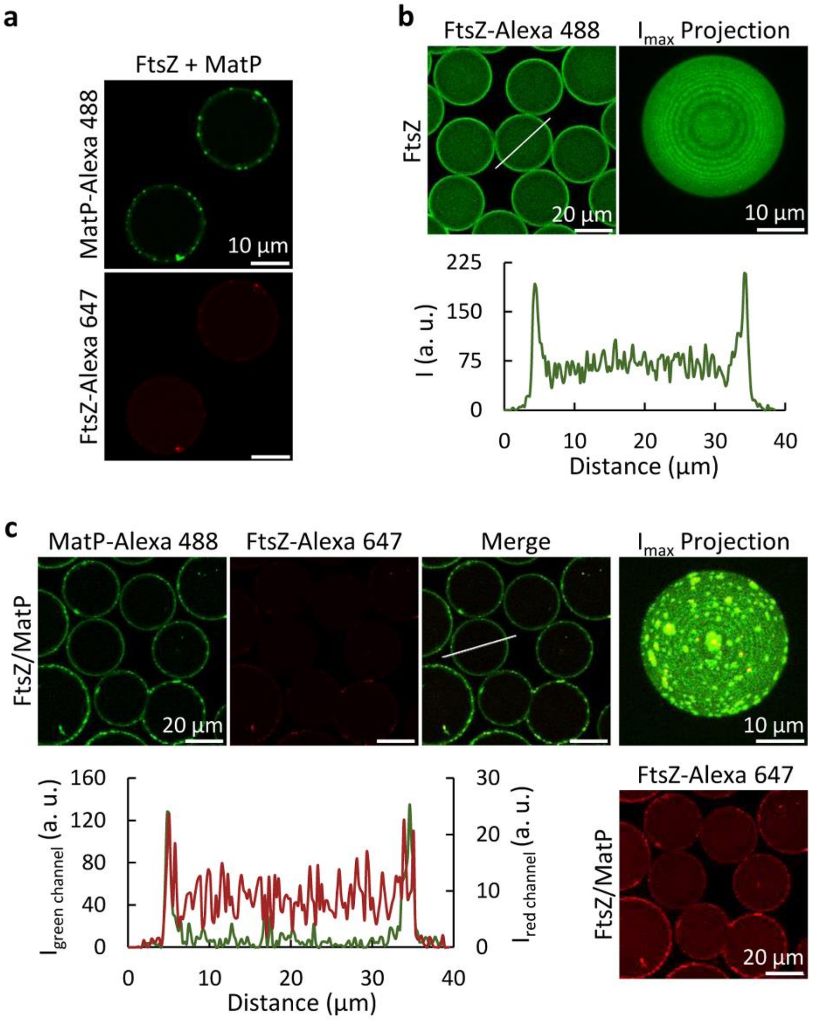


**Figure S12.** FtsZ and MatP encapsulated inside microfluidic droplets form condensates that preferentially localize at the lipid surface.
(**a**) Confocal images of the equatorial sections of microdroplets containing FtsZ and MatP mixed just before encapsulation (merge and maximum intensity projection in Figure 3f).
(**b**) Images of the equatorial section (left) and maximum intensity projection (right) of microdroplets containing FtsZ. Below, intensity profile along the line drawn in the image.
(**c**) Images of equatorial section and, on the far right, the maximum intensity projection of microdroplets containing FtsZ/MatP condensates preformed before encapsulation. Image below is that of the red channel above with a 70% increased brightness (see Materials and Methods in the main text). Intensity profiles of the green and red channels along the line depicted in the raw merge image are also shown.

Concentrations were 5 µM FtsZ, 3 µM MatP and 1 µM labeled components. Experiments were performed in *FtsZ/MatP-crowding conditions*.

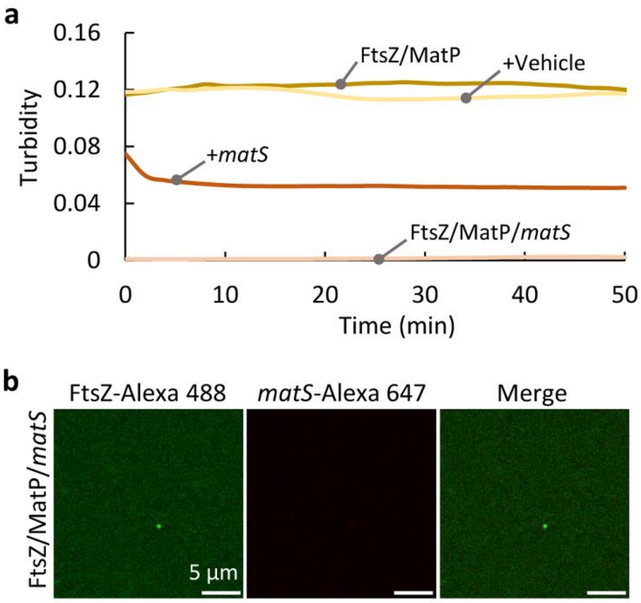


**Figure S13.** FtsZ/MatP condensates are modulated by *matS*.
(**a**) Evolution with time of the turbidity of FtsZ/MatP condensates without and with added (+) or incubated (/) *matS*. Dilution effects were discarded by adding vehicle.
(**b**) The presence of *matS* from the beginning precludes formation of FtsZ/MatP condensates.
Concentrations were 5 µM FtsZ, 3 µM MatP and 1 µM *matS* and labeled components. Experiments were performed in *FtsZ/MatP-crowding conditions*.

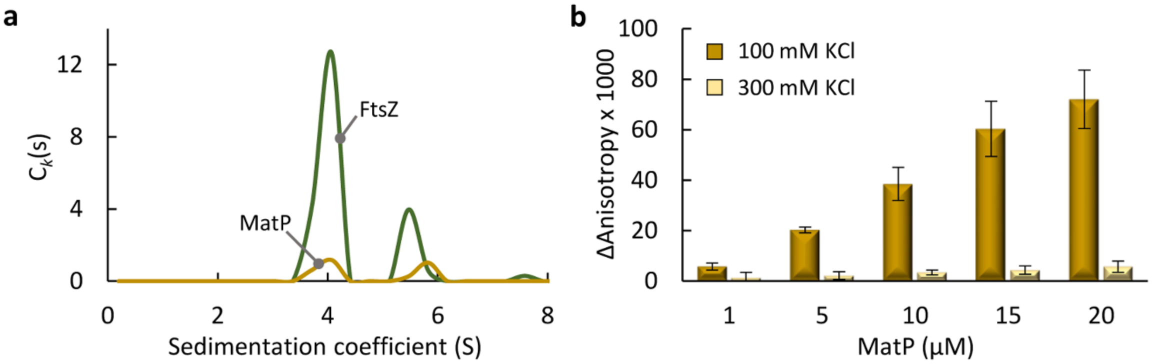


**Figure S14.** FtsZ and MatP interact in diluted solution.
(**a**) Decomposition of component sedimentation coefficient distributions, $c_k$(s), for MatP and FtsZ, obtained by global analysis of sedimentation coefficient distributions from a MSSV assay with a mixture of 20 µM FtsZ and 7 µM MatP, at 250 and 280 nm. The coexistence of both proteins within the peaks sedimenting between 4 S and 5.5 S is shown. Note that a large fraction (~50%) in these FtsZ/MatP samples consisted of higher order complexes that fully sedimented before reaching the final speed of the assay (Figure 5b).
(**b**) Change in the fluorescence anisotropy of FtsZ upon addition of MatP at the specified KCl concentrations (*n* = 4, except for 15 µM MatP at 100 mM KCl sample, *n* = 3). FtsZ concentration was 5 µM (10 nM FtsZ-Alexa 488 used as tracer). Errors are s.d.
Experiments were performed in *dilute solution buffer* with 100 mM KCl (unless otherwise stated).

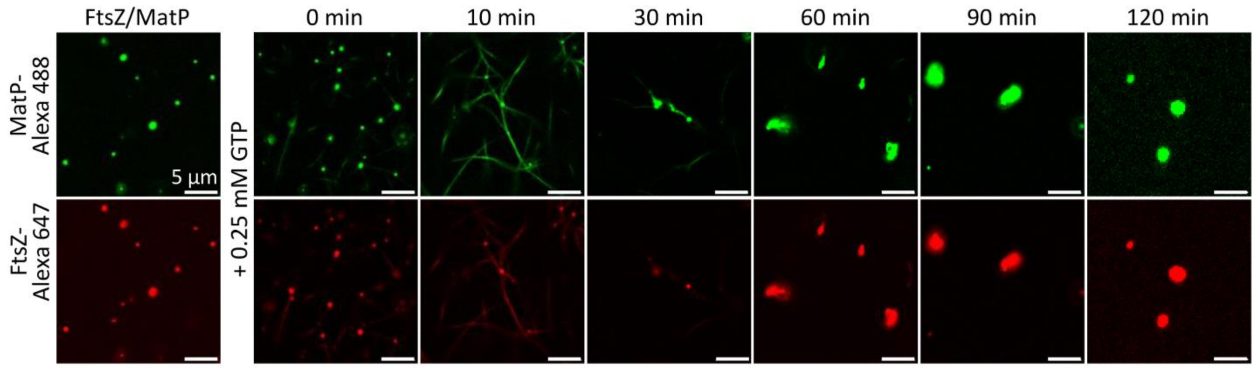


**Figure S15.** MatP decorates GTP-triggered FtsZ bundles from FtsZ/MatP condensates.
Confocal images of FtsZ/MatP condensates before and after GTP addition and time-evolution showing condensate reassembly upon GTP depletion (merge in Figure 6a). Experiments were performed in *FtsZ/MatP-crowding conditions*. Concentrations of FtsZ, MatP and labeled components were 5, 3 and 1 µM, respectively.

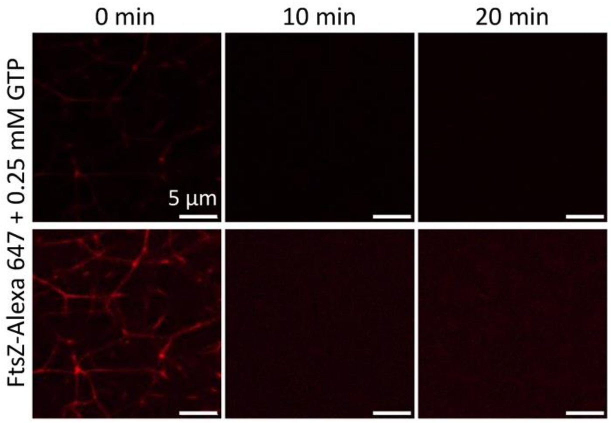


**Figure S16.** GTP-triggered FtsZ bundles depolymerization.
Confocal images of the evolution of FtsZ bundles visualized right after GTP addition (0 min) at the specified times. Images in the bottom row correspond to those above with a 50% brightness enhancement (see Materials and Methods in the main text). Concentration was 5 µM FtsZ (with 1 µM FtsZ-Alexa 647 as tracer). Experiments were performed in *FtsZ/MatP-crowding conditions*.

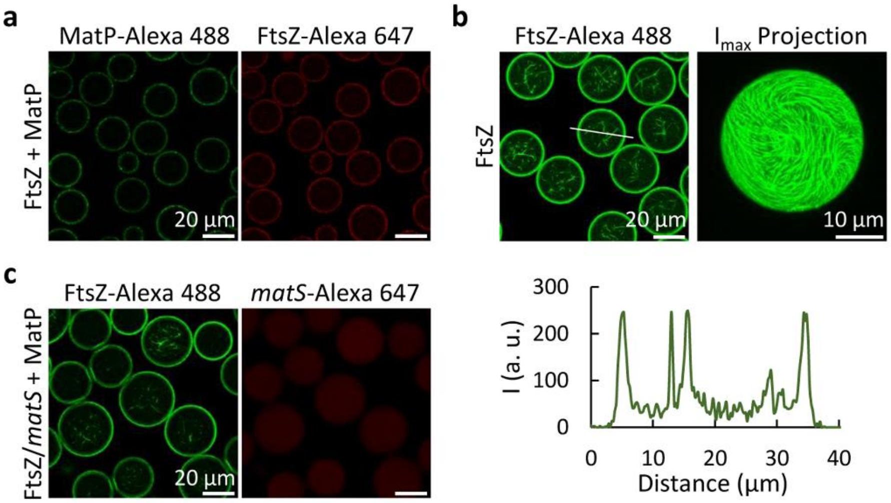


**Figure S17.** The distribution of encapsulated GTP-triggered FtsZ polymers is modified by MatP, and *matS* reverts the effect.

(**a**) Confocal images of the equatorial section of microdroplets containing FtsZ polymers and MatP (maximum intensity projections and intensity profiles in Figure 6e). FtsZ and MatP met at the droplet formation junction.

(**b**) Images of equatorial section of microdroplets containing FtsZ polymers and maximum intensity projection of a microdroplet. The intensity profile obtained along the line drawn in the image is also shown.

(**c**) Images of the equatorial section of microdroplets containing FtsZ polymers, MatP and *matS* (maximum intensity projections and intensity profiles in Figure 6f). FtsZ/*matS* and MatP met at the droplet formation junction.

Concentrations were 5 µM FtsZ, 3 µM MatP and 1 µM *matS* and labeled components. Polymerization was triggered with 2 mM GTP just before encapsulation. Encapsulations were performed in *FtsZ/MatP-crowding conditions*.

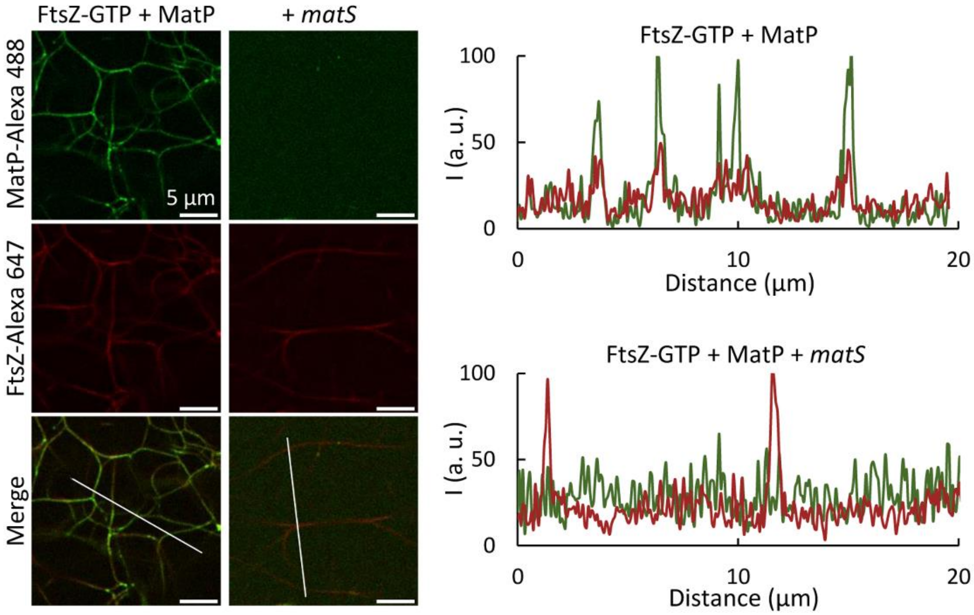


**Figure S18.** Addition of *matS* dislodges MatP from GTP-triggered FtsZ bundles.

Confocal images of FtsZ polymers (induced with 2 mM GTP) to which MatP and *matS* were sequentially added. On the right, intensity profiles obtained along the lines drawn on the images.

Concentrations were 5 µM FtsZ, 3 µM MatP and 1 µM *matS* and labeled components. Experiments were performed in *FtsZ/MatP-crowding conditions*.

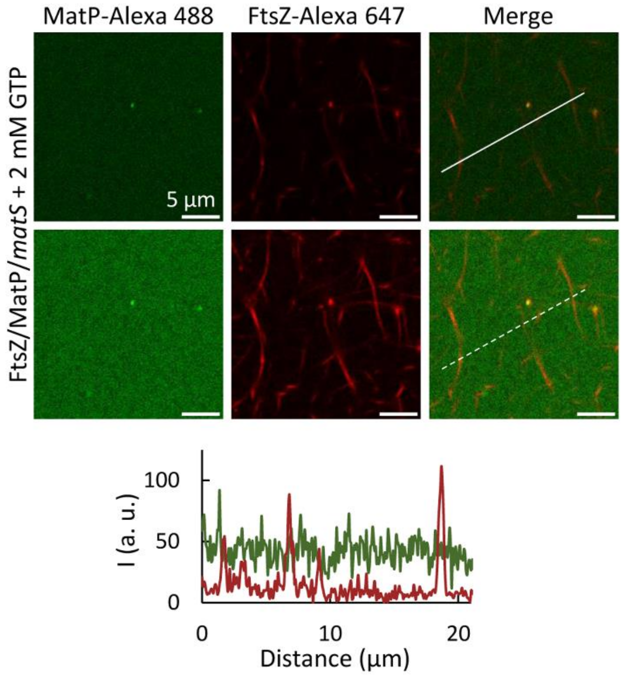


**Figure S19.** Incubation with *matS* reduces MatP incorporation into GTP-triggered FtsZ polymers.
Confocal images of samples containing FtsZ/MatP/*matS* (5, 3 and 1 µM, respectively) after addition of GTP. Images in the bottom row are those above with a 40% brightness increase (see Materials and Methods in the main text). Intensity profiles along the line drawn on the raw image (indicated as dashed line in the brightness-increased image to facilitate visualization) are shown below. Labeled components were at 1 µM. Samples were prepared in *FtsZ/MatP-crowding conditions*.

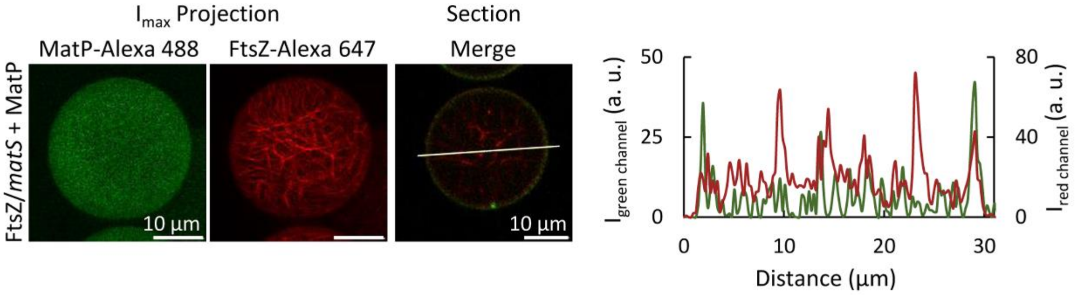


**Figure S20.** Microdroplets containing GTP-triggered FtsZ polymers, MatP and *matS.*
Maximum intensity projections and equatorial section of FtsZ polymers encapsulated with MatP and *matS* inside microdroplets. FtsZ/*matS* and MatP met at the droplet formation junction. Intensity profiles correspond to the line drawn on the image. Polymerization was triggered with 2 mM GTP right before encapsulation. The concentrations were 5 µM FtsZ, 3 µM MatP and 1 µM *matS* and labeled components. Encapsulation was performed in *FtsZ/MatP-crowding conditions*.

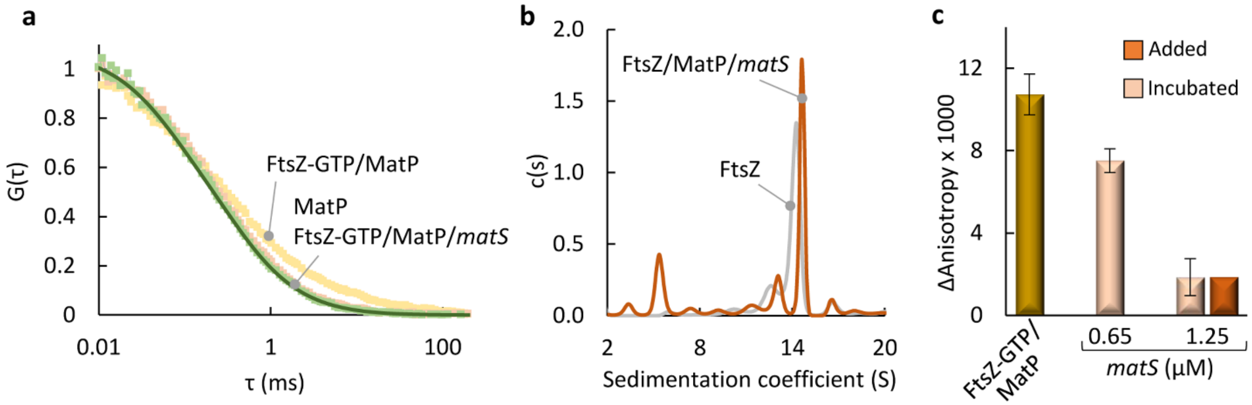


**Figure S21.** MatP interacts with GTP-triggered FtsZ polymers and *matS* disrupts these complexes.
(**a**) Normalized FCS autocorrelation curves of MatP (10 nM MatP-Alexa 488 as tracer), in the absence and presence of FtsZ and with or without *matS* (*n* > 5). Solid line is the fit of the model indicated in the main text.
(**b**) SV interference distribution of FtsZ polymers with or without MatP/*matS*. FtsZ/MatP samples contained very large complexes that substantially sedimented before reaching the final speed of the assay, in addition to other species that remained in solution; evolution of this sample during the experiment precluded quantitative analysis to obtain sedimentation coefficient distributions.
(**c**) Change of the fluorescence anisotropy of FtsZ polymers (50 nM FtsZ-Alexa 488 as tracer) with MatP, without or with *matS*, incubated with the proteins before triggering polymerization or added right after polymerization of FtsZ/MatP with GTP. Values are the average of 3 independent experiments ± s.d.
Concentrations were 10 (a), 5 (b) or 2.5 μM (c) MatP; 20 (a), 10 (b) or 5 μM (c) FtsZ; 5 (a), 2.5 μM (b) or as specified (c) *matS*. Polymerization was triggered by addition of 2 mM GTP with (a, b) or without RS (c). Experiments were conducted in *dilute solution buffer* with 100 mM KCl.

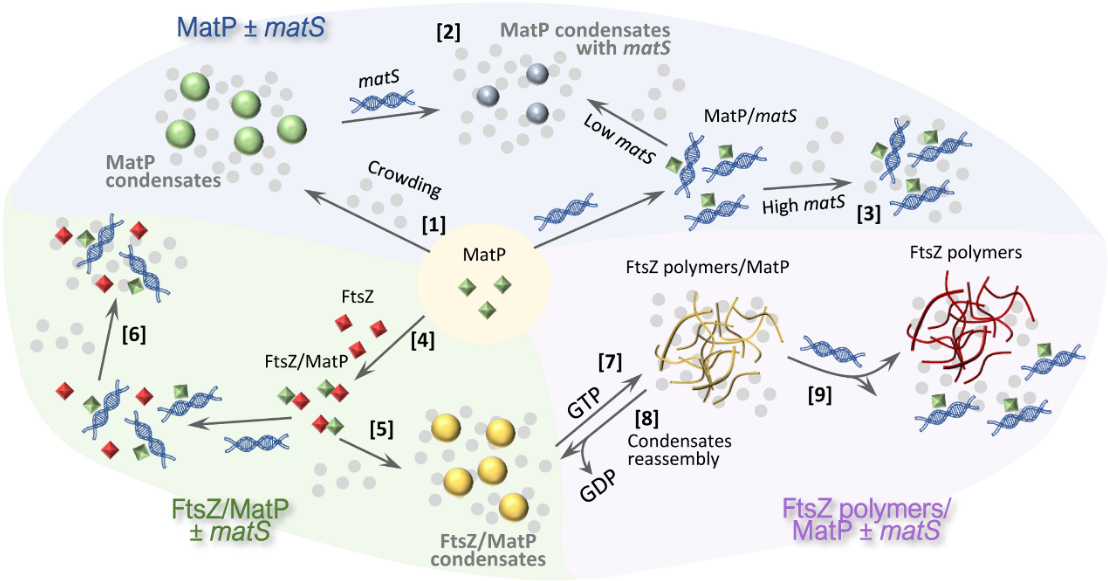


**Figure S22.** Schematic representation of the assembly and regulation of MatP biomolecular condensates.

MatP forms crowding-driven condensates [1] where *matS* can incorporate [2], although high *matS* concentration precludes phase separation [3]. MatP interacts with FtsZ [4] and forms heterotypic condensates in crowding conditions [5]. *matS* disrupts FtsZ/MatP complexes, hindering condensation [6]. GTP triggers FtsZ polymers with bound MatP from FtsZ/MatP condensates [7] that reassemble after GTP depletion [8]. *matS* dissociates MatP from FtsZ polymers [9].

### Legends for Supplementary movies

**Movie 1.** 3D projections of GTP-triggered FtsZ polymers with MatP encapsulated inside a microfluidic droplet stabilized by lipids, as described in Figure 6e. Bottom, 3D projections without interpolation.

**Movie 2.** 3D projection of a lipid-stabilized microfluidic droplet containing GTP-triggered FtsZ polymers. On the right, 3D projection without interpolation. Encapsulation was conducted under the conditions specified in Figure S17b.

**Movie 3.** 3D projections of a lipid-stabilized microfluidic droplet containing GTP-triggered FtsZ polymers, MatP and *matS*. Bottom, 3D projections without interpolation. Encapsulation was conducted under the conditions described in Figure S20.

### Supplementary Materials and Methods

#### Reagents

Polar extract phospholipids from *E. coli* were from Avanti Polar Lipids (AL, USA), and stored in chloroform at −20 °C until used. GTP and dextran 500 (500 kDa) were from Merck (NJ, USA). Ficoll 70 (70 kDa) was from GE Healthcare (IL, USA). The concentrations of dextran and Ficoll were measured from the refractive index increment as described (1). Alexa Fluor 488 and Alexa Fluor 647 carboxylic acid succinimidyl ester dyes were from Thermo Fisher Scientific Inc. (MA, USA). Analytical-grade chemicals were from Merck. Single stranded oligonucleotides purified by HPLC containing the *matS19* sequence (AAA<u>GTGACACTGTCAC</u>CTT, bases recognized by MatP underlined (2)) or a nonspecific sequence of similar length (AAGTAAGTGAGCGCTCACTTACGT), either unlabeled or labeled with Alexa Fluor 647 or fluorescein in the 5′ end, were obtained from IDT. BSA (Merck) was dialyzed in 50 mM Tris-HCl pH 7.5, 300 mM KCl. The enzymatic GTP regeneration system (RS) contained 2 units/mL acetate kinase and 15 mM acetyl phosphate (3), both from Merck.

#### Confocal microscopy

Samples, either crowding bulk solutions or microfluidic droplets, were visualized in silicone chambers (Thermo Fisher Scientific) glued to coverslips, using a Leica TCS SP5 inverted confocal microscope with a HCX PL APO 63× oil immersion objective (N.A. = 1.4; Leica, Mannheim, Germany). Excitation of Alexa Fluor 488 and Alexa Fluor 647 was with 488 nm and 633 nm laser lines, respectively. Each individual sample was analyzed by registering several images at different observation fields. Samples with condensates were incubated for 30 min before visualization and, when monitoring the effect of additional elements (proteins or DNA) directly added over the sample, they were imaged before and immediately after the addition or at the specified time. Evolution of polymers with time was monitored by preparing several samples with the same composition and adding, after the 30 min incubation, GTP at different times.

### Fluorescence anisotropy

Anisotropy experiments were performed in a Spark® Multimode microplate reader (Tecan) with 485 and 535 nm excitation and emission filters, respectively. Samples were prepared in Protein LoBind tubes and transferred to 384-wells black polystyrene (non-binding surface), flat bottom microplates (Corning). The temperature was regulated at 26 °C. FtsZ/MatP binding isotherms were obtained using 50 nM MatP-Alexa 488 as tracer, in the same buffer used for the FCS experiments, with 100 mM KCl. Reported anisotropy values are the average of three independent replicates ± s.d. Binding analysis was conducted using BIOEQS software (4, 5), with a simple 1:1 model compatible with the data, to obtain apparent *Kd*s corresponding to the concentration of FtsZ at which half of maximum signal is observed, under our experimental conditions. This analysis takes into account the concentration of MatP. An increase in the total fluorescence intensity occurred as the concentration of FtsZ increased, not due to any background signal from FtsZ (background fluorescence was subtracted in all samples). Quenching of Alexa Fluor 488 fluorescence by amino acids, described in previous reports (6), might occur in MatP-Alexa 488 and be reverted by interaction with FtsZ. However, fluorescence lifetime measurements, using the Microtime 200 instrument described in the main text, showed no significant difference in the average lifetime (around 2.5-2.6 ns) between free MatP and FtsZ/MatP complexes, ruling out dynamic quenching. Since the quenching of Alexa Fluor 488 by amino acids usually results from either dynamic or both dynamic and static quenching (6), the lack of dynamic quenching proves this option very unlikely. The increase in intensity could be explained, instead, by a reduction of the high tendency of MatP-Alexa 488 to adsorb to surfaces upon increasing FtsZ concentration, which would not affect the average anisotropy at each FtsZ concentration nor the apparent binding constants, well above the MatP-Alexa 488 concentration. Nevertheless, BIOEQS analyses were also conducted considering FtsZ/MatP complexes with larger intensity than free MatP, rendering binding parameters only slightly different than those in which equal intensity contribution was assumed (apparent $\Delta G$ 6.5 ± 0.2 *vs* 7.0 ± 0.2 kcal/mol; apparent $K_d$ 18 ± 6 *vs* 8 ± 3 μM). Uncertainties in the parameters retrieved were calculated with the same software by rigorous confidence limit testing at the 67% level, and error propagation in the case of the apparent $K_d$.
